# Pparα and fatty acid oxidation coordinate hepatic transcriptional architecture

**DOI:** 10.1101/2021.07.12.451949

**Authors:** Kyle S. Cavagnini, Michael J. Wolfgang

**Author notes:** Address correspondence to: **Michael J. Wolfgang, Ph.D.**, Department of Biological Chemistry, Johns Hopkins University School of Medicine, 855 N. Wolfe St., 475 Rangos Building, Baltimore, MD 21205.

## Abstract

Fasting requires tight coordination between the metabolism and transcriptional output of hepatocytes to maintain systemic glucose and lipid homeostasis. Genetically-defined deficits in hepatic fatty acid oxidation result in dramatic fasting-induced hepatocyte lipid accumulation and induction of genes for oxidative metabolism, thereby providing a mouse model to interrogate the mechanisms by which the liver senses and transcriptionally responds to fluctuations in lipid levels. While fatty acid oxidation is required for a rise in acetyl-CoA and subsequent lysine acetylation following a fast, changes in histone acetylation (total, H3K9ac, and H3K27ac) associated with transcription do not require fatty acid oxidation. Instead, excess fatty acids prompt induction of lipid catabolic genes largely via ligand-activated Pparα. We observe that active enhancers in fasting mice are enriched for Pparα binding motifs, and that inhibition of hepatic fatty acid oxidation results in elevated enhancer priming and acetylation proximal to Pparα binding sites within regulatory elements largely associated with genes in lipid metabolism. Also, a greater number of Pparα-associated H3K27ac signal changes occur at active enhancers compared to promoters, suggesting a genomic mechanism for Pparα to tune target gene expression levels. Overall, these data demonstrate the requirement for Pparα activation in maintaining transcriptionally permissive hepatic genomic architecture particularly when fatty acid oxidation is limiting.

**HIGHLIGHTS:**

- Fasting-induced transcription and histone acetylation are largely independent of acetyl-CoA concentration.
- Deficits in fatty acid oxidation prompt epigenetic changes and Pparα-sensitive transcription.
- Fasting prompts enhancer priming and acetylation proximal to Pparα binding sites independent of Pparα.
- Patterns of Pparα target genes can be distinguished by epigenetic marks at promoters and enhancers.

## INTRODUCTION

The liver is a principle regulator of systemic lipid physiology. In this role the hepatocyte requires signaling mechanisms by which to sense and respond to fluctuations in lipid availability. This is especially important during periods of nutrient deprivation, such as fasting (George, 2006). Fasting stimulates fatty acid mobilization from adipose, whereby they are taken up and broken down in the liver via mitochondrial β-oxidation to provide hepatocytes with ATP, NADH, and acetyl-CoA (Stern et al., 2016; The and Schulz, 1991). Errors in these fundamental catabolic processes result in metabolic aberrations, such as fasting hypoketotic-hypoglycemia, and can also contribute to the pathogenesis and progression of conditions such as diabetes, obesity, and chronic liver disease (Asrani et al., 2019; Gong et al., 2017; Houten et al., 2016; Ponziani et al., 2015).

The nuclear hormone receptor peroxisome proliferator-activated receptor alpha (Pparα) plays a governing role in regulating hepatic lipid homeostasis and is thus a key component of the fasting response (Kersten, 2014; Kersten et al., 1999; Leone et al., 1999). Upon activation by lipid ligands, such as long chain fatty acids, Pparα and its heterodimer partner retinoid X receptor alpha (RXRα) will bind to DNA and effect transcription of target genes (Bardot et al., 1993; Boergesen et al., 2012; Evans and Mangelsdorf, 2014; Gearing et al., 1993). This transcriptional program includes genes for mitochondrial and peroxisomal β-oxidation, microsomal ω-oxidation, and ketogenesis (Aoyama et al., 1998; Mandard et al., 2004; Rakhshandehroo et al., 2009). It prompts induction of regulators for mitochondrial metabolism such as carnitine palmityltransferase 1a (*Cpt1a*) and pyruvate dehydrogenase kinase 4 (*Pdk4*), which respectively promote mitochondrial fatty acid import and inhibit pyruvate oxidation, (Huang et al., 2002; Song et al., 2010; Wu et al., 2001). Pparα also activates transcription of pro-catabolic hepatokines such as fibroblast growth factor 21 (*Fgf21*), which help mediate the adaptive starvation response systemically (Inagaki et al., 2007; Iroz et al., 2017). Pparα knockout mice display diminished transcription of these and other genes for oxidative metabolism, which over time contributes to hepatic lipid accumulation and steatosis (Aoyama et al., 1998; Kersten et al., 1999; Leone et al., 1999; Reddy, 2001; Ruppert et al., 2019).

Transcription factor binding hubs, such as promoters and enhancer elements, play a fundamental role in regulating hepatic transcription (Goldstein and Hager, 2015; Jump et al., 2013; Karagianni and Talianidis, 2015; Qin et al., 2020). Enhancers are cis-regulatory genetic elements that facilitate transcription initiation and processivity; they are classified as silenced, poised, or active (Creyghton et al., 2010). Poised enhancers are marked by enrichment of epigenetic modifications including H3K4me1 and H3K27me3 (Calo and Wysocka, 2013; Heinz et al., 2015). Active enhancers are marked by a coincidence of H3K4me1 and H3K27ac, as well as increased chromatin accessibility and the presence of histone acetyltransferases such as p300 (Andersson and Sandelin, 2020; Creyghton et al., 2010; Raisner et al., 2018). The hepatic fasting response requires careful balance between gluconeogenesis and ketogenesis. The interplay between each process is in part governed by transcription factor binding patterns, including changes in the active enhancer landscape. Glucocorticoid receptor (GR)-assisted loading of cAMP responsive element binding protein I promotes activation of gluconeogenic enhancers (Goldstein et al., 2017). In contrast, ketogenic enhancers are proposed to have a more gradual enhancer maturation which is correlated to GR-stimulated expression of Pparα and increased chromatin accessibility in proximity to Pparα binding motifs (Goldstein et al., 2017). Moreover, Pparα binding has been directly detected at fasting-induced enhancers using ChIP-seq (Lee et al., 2014; Sommars et al., 2019). Pparα-null mice have suppressed indicators for active enhancers, including decreased enhancer RNA levels and H3K27ac ChIP-seq signal (Guan et al., 2018; Sommars et al., 2019).

In this study we take advantage of mice with a genetically-defined deficit in hepatic fatty acid oxidation to further investigate the contribution of Pparα in regulating the hepatic fasting transcriptional landscape, including a close inspection of its activity at active enhancers. Carnitine palmityltransferase 2 (Cpt2) is a required enzyme for the transport of long chain fatty acids into the mitochondria (Houten et al., 2016). Its loss deprives the mitochondria of fatty acid substrate, and it is thus an obligate enzyme for mitochondrial β-oxidation. During a fast, mice with a liver-specific deletion of Cpt2 (Cpt2^L−/−^) are unable to clear excess lipid (Lee et al., 2016). This results in drastic fatty acid accumulation accompanied by augmented transcription of many Pparα target genes (Lee et al., 2016). These mice are therefore a model of Pparα activation by build-up of endogenous lipid ligands. We use this system to further explore the effect of hepatic lipid sensing on gene activation, and the role for Pparα activation in maintaining transcriptionally permissive chromatin. In particular we describe distinct patterns of Pparα target gene transcription that are distinguished by epigenetic marks at promoters and enhancers. We further observe that Pparα-sensitive enhancers are largely associated with lipid metabolism, and deficits in fatty acid oxidation prompt elevated priming and acetylation at these loci. Altogether our findings provide genomic insight into how fatty acids alter the epigenome to affect transcription during periods of nutrient deprivation.

## RESULTS

### Impaired hepatic mitochondrial fatty acid oxidation elicits putative Pparα target genes despite a suppression in lysine acetylation

Mice lacking hepatic Cpt2 (Cpt2^L−/−^) are unable to utilize long chain fatty acids for mitochondrial β-oxidation. Following a fast, this results in fatty liver and dramatic induction of pro-catabolic hepatic genes, thereby providing a useful genetic model to interrogate fatty-acid stimulated transcription (Bowman et al., 2019; Lee et al., 2016). To fully characterize the transcriptional landscape of the fasting Cpt2^L−/−^ liver, we performed RNA-seq on livers harvested from 9-week old wildtype (WT) and Cpt2^L−/−^ mice fasted for 24 hours (**(Fig. 1a, Table S1**). This expanded upon our previously published observations that impaired hepatic β-oxidation is associated with augmented fasting-induced transcription of genes for regulation of mitochondrial metabolism (*Pdk4*), regulation of peroxisomal metabolism (*Ehhadh*), and pro-catabolic hepatokines (*Fgf21*) (Lee et al., 2016). Indeed, gene ontology for all significantly upregulated transcripts (Cpt2^L−/−^/WT fold change ≥ 2, padj < 0.05) returned several terms for fatty acid metabolism (**Fig. S1a**). We next performed TMT-based quantitative proteomics to confirm the induction of these genes at the protein level (**Table S2**). We found that, in parallel to the RNA-seq data, Cpt2^L−/−^ upregulated peptides were enriched for gene ontology terms related to fatty acid biology (**Fig. S1b, S1c**). These results provide additional evidence demonstrating that impaired β-oxidation triggers a compensatory transcriptional program in an attempt to relieve the lipid burden, a process which includes shuttling fatty acids into other metabolic pathways such as peroxisomal oxidation.

**Fig 1.**
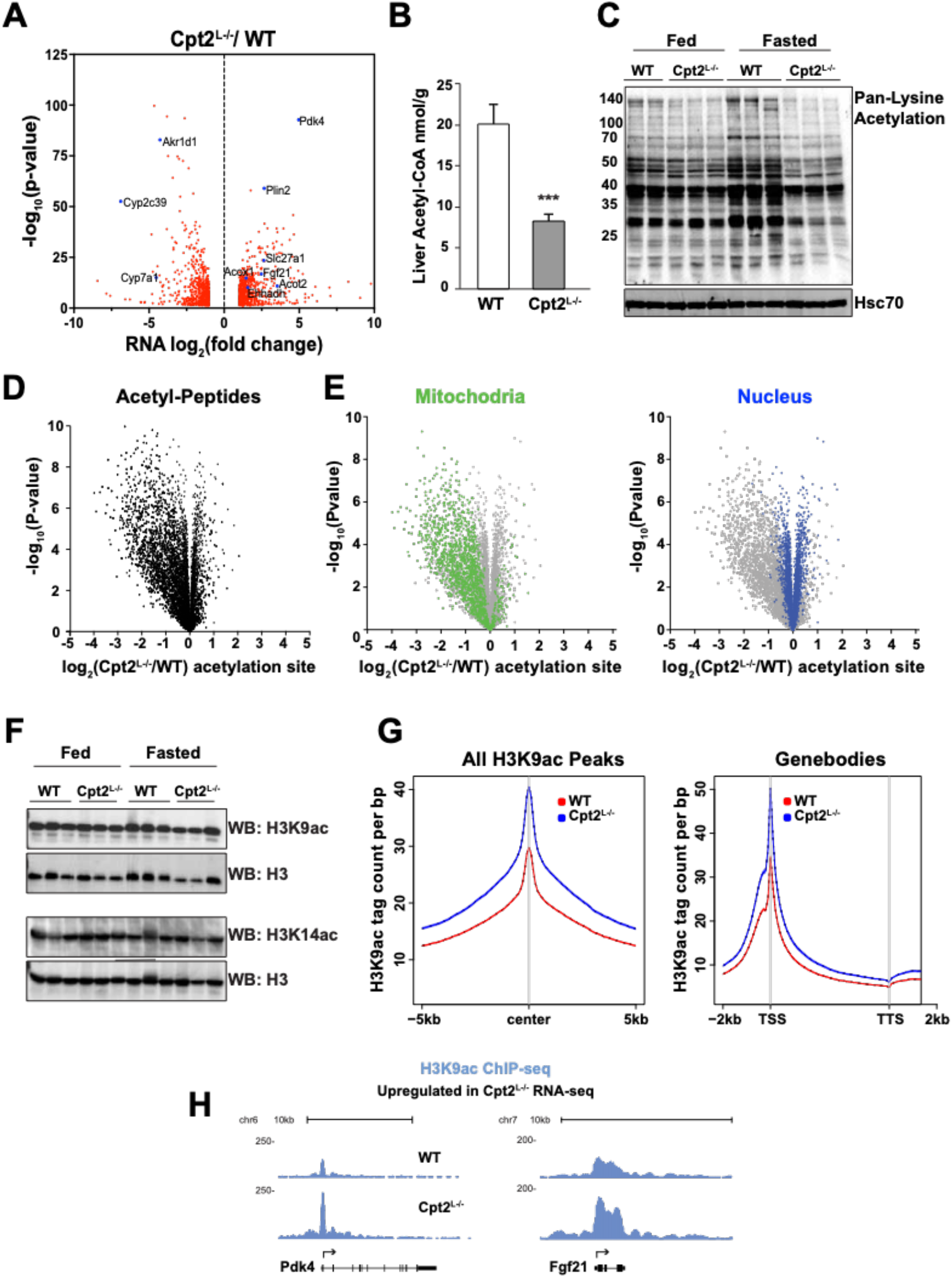
Fasted Cpt2^L−/−^ liver exhibits deficits in lysine-acetylation along with metabolic gene induction. A. Volcano plot displaying differentially expressed transcripts (fold change ≥ |2|, padj < 0.05) between 24hr fasted WT and Cpt2^L−/−^ liver as determined by RNA-seq (n=4). B. Tissue acetyl-CoA concentration from 24hr fasted WT and Cpt2^L−/−^ liver (n=6, mean ± SEM). Significance determined using Student two-tailed t test, ***p < 0.001. C. Western blot for acetyl-lysine and HSC70 loading control in fed and fasted WT and Cpt2^L−/−^ livers. D. Volcano plots showing magnitude and significance for fasted liver acetyl-proteome, measured by TMT-based quantitative mass spectrometry (n=5). E. Volcano plots showing magnitude and significance for mitochondrial acetyl-proteome and nuclear acetyl-proteome. F. Representative Western blot of acid-precipitated histones from fed and fasted WT and Cpt2^L−/−^ liver tissue. G. Aggregation plot depicting liver H3K9ac ChIP-seq mean tag density for (*left*) all H3K9ac ChIP-seq peak centers with ±5kb flanking regions and (*right*) across all gene bodies with ±2kb flanking regions. TSS = transcription start site, TTS = transcription termination site. n=1 H. H3K9ac ChIP-seq genome browser tracks for representative genes that are upregulated in Cpt2^L−/−^ RNA-seq, showing increased ChIP-seq signal in the fasted Cpt2^L−/−^ liver.

Others have suggested that fatty acid oxidation promotes transcription for lipid catabolism via histone acetylation from fatty acid-derived acetyl-CoA (McDonnell et al., 2016). We observed that Cpt2^L−/−^ mice exhibited suppressed hepatic acetyl-CoA concentration and failed to induce protein acetylation following a fast, which therefore allowed us to examine this hypothesis *in vivo* (**Fig. 1b, 1c**). To gain a more granular view of the proteins that exhibited hypoacetylation, we mapped and quantified protein acetylation via TMT-based mass spectrometry (**Fig. 1d, Table S3**). We found that mitochondrial protein acetylation was suppressed in Cpt2^L−/−^ mice as expected due to a lack of mitochondrial β-oxidation (**Fig. 1e, S1e**). In fact, lysine acetylation was globally reduced in Cpt2^L−/−^ mice with the noted exception of nuclear peptides, which retained comparable levels of acetylation between WT and Cpt2^L−/−^ mice (**Fig. 1e, S1d**). These data indicate that the deficit in mitochondrial β-oxidation limits liver acetyl-CoA substrate for lysine acetylation, with the exception of the nucleus where acetyl-lysine levels appear buffered.

Analysis of acetyl-histone peptides provides evidence against bulk histone acetylation as a primary driver of transcriptional activation. 91 unique acetyl-histone peptides were detected in the acetyl-proteome, of which none were hyperacetylated, 38.5% were hypoacetylated and the remaining 61.5% showed no significant change to acetylation state between Cpt2^L−/−^ and WT animals (**Table S3**). Consistent with those data, bulk changes in histone acetylation were not observed for histone marks H3K9ac or H3K14ac (**Fig. 1f**). To gain detailed insight into fatty acid oxidation-dependent histone acetylation, we next turned to ChIP-seq for H3K9ac. Global H3K9ac levels were higher in Cpt2^L−/−^ liver, including at gene promoters, consistent with the augmented transcriptional profile of Cpt2^L−/−^ mice (**Fig. 1g**). Local patterns of H3K9ac gene body occupancy trended with RNA-seq fold-change (**Fig. 1h, S1f**). These data are not consistent with the notion that the fatty acid oxidation-dependent generation of acetyl-CoA is a primary driver of gene expression via an epigenetic mechanism, but rather point towards another lipid sensing mechanism for transcriptional activation.

To better understand how fatty acids affect the chromatin landscape, including promotor accessibility and binding motifs for transcription factors induced in fasting Cpt2^L−/−^ liver, we next turned to Assay for Transposase Accessible Chromatin Sequencing (ATAC-seq). ATAC-seq profiles of fasted Cpt2^L−/−^ and WT liver tissues were overall similar. A total of 69,485 ATAC-seq sites were measured across all samples, of which 75.6% are shared by both fasted Cpt2^L−/−^ and WT animals (**Fig. 2a**). Cpt2^L−/−^ livers had a notable increase in chromatin accessibility near transcription start sites (**Fig. 2b**). Of the 9273 differentially upregulated ATAC sites (fold change Cpt2^L−/−^/WT ≥ +2), 16% were detected in gene promotors. Differential accessibility aligned with RNA-seq trends (**Fig. 2c, S2a**). Motif analysis on ATAC peaks found at gene promotors for upregulated Cpt2^L−/−^ transcripts revealed Pparα and its heterodimer binding partner Rxr as the top hits (**Fig. 2d**). Pparα targets are often defined by their response to a Pparα agonist, such as WY-14643 (Brocker et al., 2020; Janssen et al., 2015; Lee et al., 1995; Li et al., 2018; Rakhshandehroo et al., 2007). We determined that 52% of genes upregulated in Cpt2^L−/−^ animals are sensitive to WY-14643 (**Fig. 2e**) (Naiman et al., 2019). Together, the proteomic, acetyl-proteomic, RNA-seq, ATAC-seq and H3K9ac ChIP-seq data suggest that deficits in hepatic fatty acid oxidation by the loss of Cpt2 elevates endogenous lipid ligands to induce Pparα at promoters of genes important for lipid catabolism despite a suppression in acetyl-CoA.

**Fig 2.**
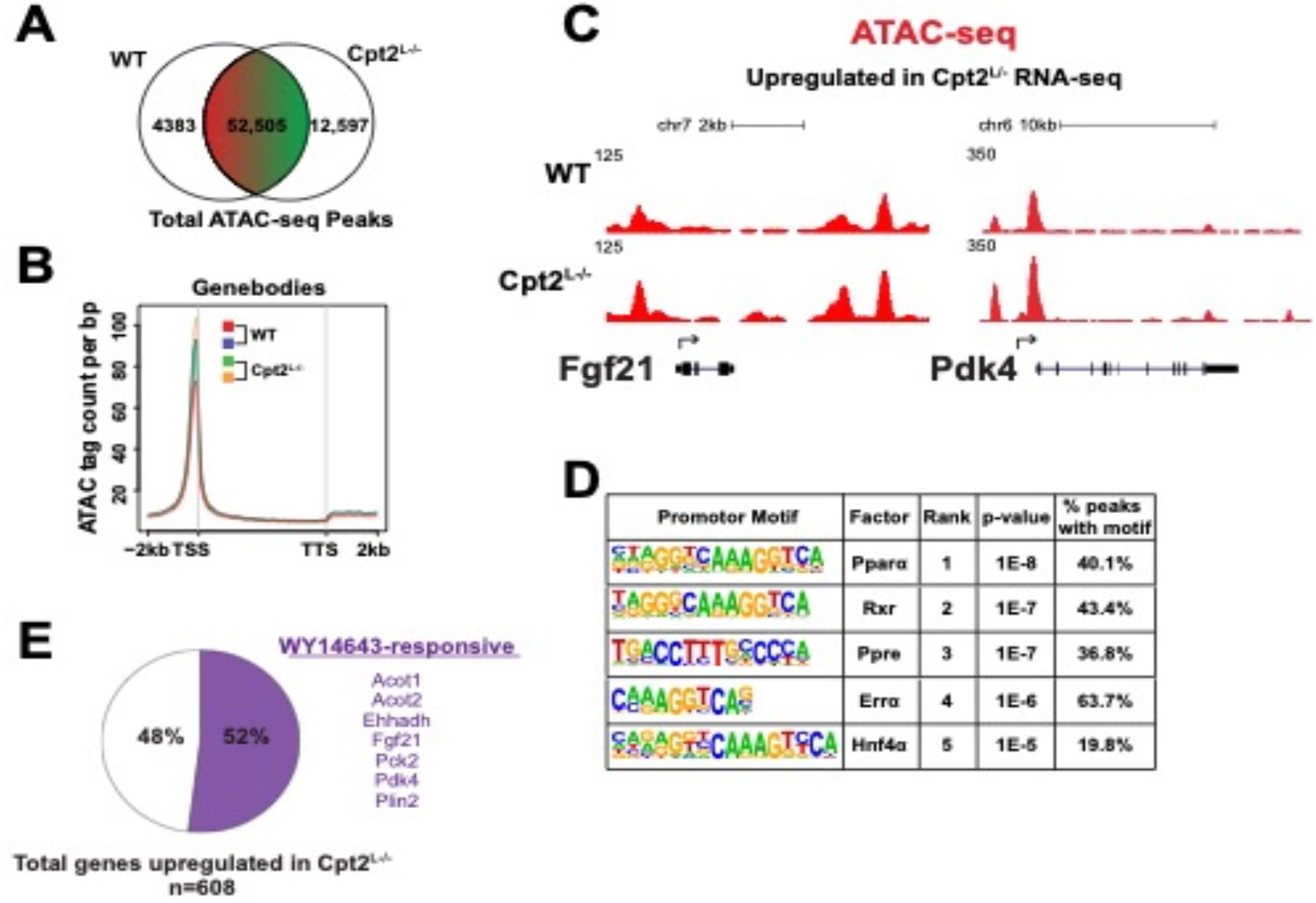
Fasting-induced genes in Cpt2^L−/−^ liver are Pparα-responsive. A. Venn diagram showing unique and shared ATAC-seq peaks detected in fasted WT and Cpt2^L−/−^ liver (n=2). B. Aggregation plot depicting liver ATAC-seq mean tag density for all gene bodies with ±2kb flanking regions. TSS = transcription start site, TTS = transcription termination site. C. ATAC-seq genome browser tracks for representative genes that were upregulated in Cpt2^L−/−^ RNA-seq. D. Top results from HOMER enrichment analysis for known motifs in ATAC-seq promoter peaks for genes induced in Cpt2^L−/−^ liver. E. Pie chart depicting Cpt2^L−/−^ induced genes previously shown to respond to the Pparα agonist WY-14643 (GSE140063, Naiman et al., 2019).

### Pparα, Cpt2^L−/−^ double knockout mice demonstrate Pparα dependent and independent transcription

Given the abundance of putative Pparα target genes and enrichment for Pparα promoter binding motifs in fasted Cpt2^L−/−^ livers, we decided to define the requirement for Pparα in the transcriptional response of Cpt2^L−/−^ mice by generating double knockout (DKO) mice. Cpt2^L−/−^;Pparα^−/−^ DKO animals were generated using mice harboring a whole-body deletion of Pparα (Pparα^−/−^) bred to Cpt2^L−/−^ mice. Subsequent experiments in this study were conducted using 24hr fasted 9-week old WT, Cpt2^L−/−^, Pparα^−/−^, and DKO mice. All genotypes were viable and fertile and were not associated with changes in body weight (**Fig. 3a**). We have previously shown that fasted Cpt2^L−/−^ mice have significantly increased liver mass due to excess fatty acids; the DKO mice share this phenotype (**Fig. 3b**). While Cpt2^L−/−^ mice did not exhibit differences in fasting glycemia compared to controls, Pparα^−/−^ and DKO mice exhibited mild fasting hypoglycemia (**Fig. 3c**). This phenotype was previously described for fasted Pparα^−/−^ animals (Kersten et al., 1999). Pparα^−/−^ mice had approximately half of the circulating fasting ketone bodies such as β-hydroxybutyrate (βHB) compared to WT animals (**Fig. 3d**). However, Cpt2^L−/−^ and DKO mice did not produce βHB following a 24hr fast, demonstrating a requirement for hepatic fatty acid oxidation for ketone body generation.

**Fig 3.**
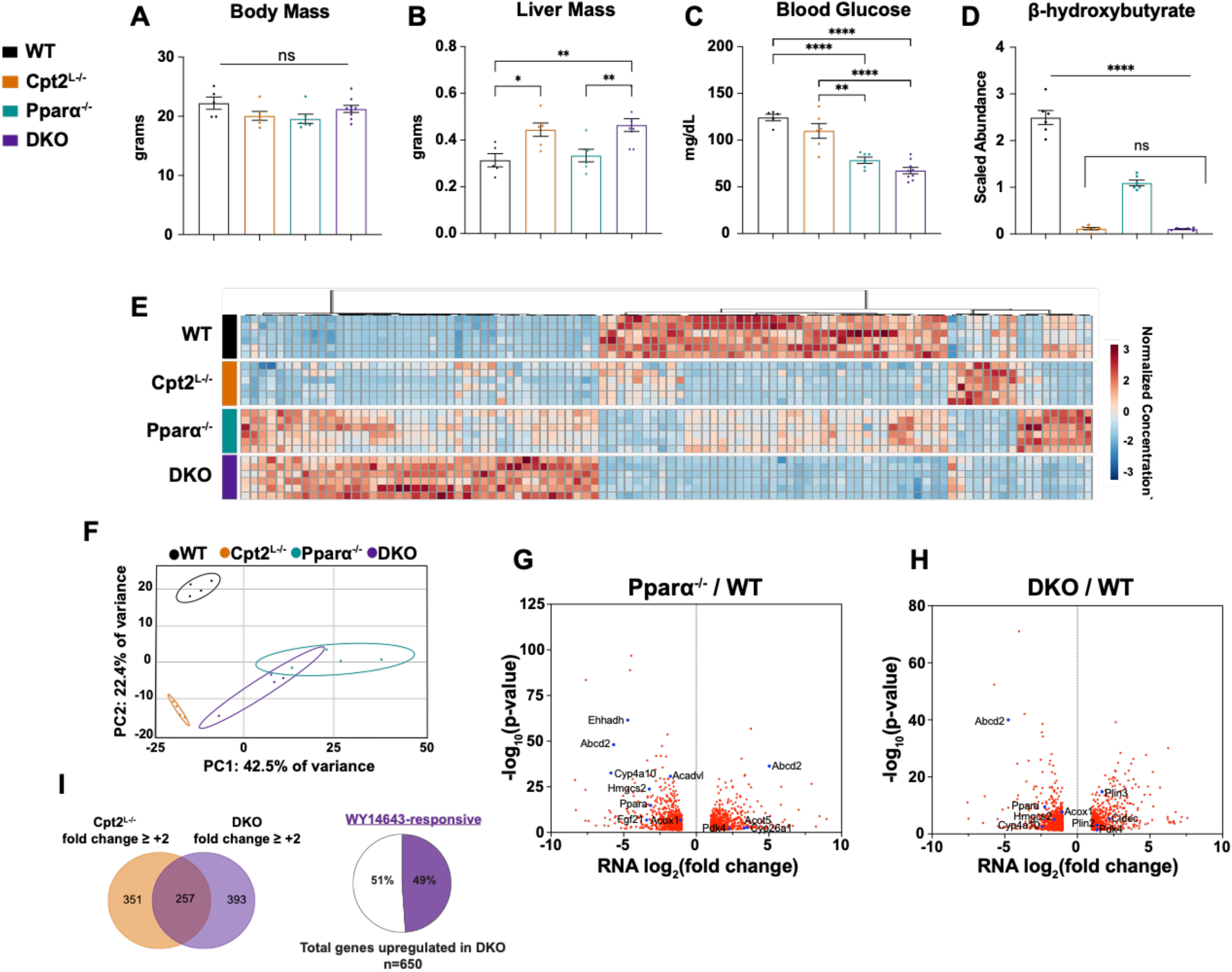
Pparα, Cpt2^L−/−^ double knockout mice retain transcription of Pparα target genes. A. Body weight of male fasted WT, Cpt2^L−/−^, Pparα^−/−^, and Cpt2^L−/−^;Pparα^−/−^ (DKO) animals (n=6-10 mean ± SEM). B. Wet liver weight from fasted animals (n=6-10, mean ± SEM). C. Blood glucose from fasted animals (n=6-10, mean ± SEM). D. Serum levels of β-hydroxybutyrate from fasted animals determined by median-scaled untargeted metabolomics (n=6, mean ± SEM). E. Heatmap of top 100 differentially regulated serum metabolites from fasted animals (n=6). F. Principle component analysis of RNA-seq on fasted WT, Cpt2^L−/−^, Pparα^−/−^, and DKO liver (n=4). G. Volcano plot displaying RNA-seq differentially expressed transcripts between 24hr fasted WT and Pparα^−/−^ livers. H. Volcano plot displaying RNA-seq differentially expressed transcripts between 24hr fasted WT and DKO livers. I. (*left*) Venn diagram depicting overlap between genes upregulated in Cpt2^L−/−^ and DKO liver compared to WT. (*right*) Pie chart depicting DKO fasting-induced genes known to respond to the Pparα agonist WY14643. RNA-seq significance cutoff is fold change ≥ |2|, padj < 0.05. One-way ANOVA followed by Tukey’s post-hoc test was performed as appropriate. *p < 0.05; **p < 0.01; ***p < 0.001; ****p < 0.0001; ns, not significant.

To better understand the physiological impact of impaired hepatic β-oxidation and Pparα loss, we performed global untargeted metabolomics on serum (**Fig. 3e, Table S4**). Principle component analysis (PCA) showed that loss of Pparα caused broad perturbations to the serum metabolome, more so than loss of Cpt2^L−/−^ alone (**Fig. 3e, S3a**). This emphasizes the importance of Pparα for systemic fasting physiology. These perturbations were compounded in the DKO animals, which showed the greatest variation from WT controls. We’ve previously reported that fasted Cpt2^L−/−^ mice show depleted serum short- and medium-chain acyl-carnitines (Lee et al., 2016). DKO mice phenocopy Cpt2^L−/−^ serum levels of medium- and long-chain acyl carnitines, indicating that this is likely a shared metabolic sink for the excess fatty acid in Cpt2^L−/−^ livers (**Fig. S3b**). These data overall demonstrate that the loss of Pparα and/or the loss of hepatic fatty acid oxidation result in systemic metabolic perturbations.

We next sought to determine the extent to which Pparα is implicated in the fasting-induced Cpt2^L−/−^ transcriptional response by carrying out RNA-seq on WT, Cpt2^L−/−^, Pparα^−/−^, and DKO livers (**Table S1**). PCA of RNA-seq data revealed that the transcriptional signature for Pparα^−/−^ mice overlapped with that of DKO mice, while those for WT and Cpt2^L−/−^ animals were distinct from the other genotypes (**Fig. 3f**). There were 330 differentially expressed transcripts shared between the three knockout genotypes compared to WT animals (KO/WT fold change ≥ |2|, padj < 0.05) (**Fig. S3c**). In line with overall transcriptional signatures, a greater number of differentially expressed genes were shared between the Pparα^−/−^ and DKO mice compared to Cpt2^L−/−^ mice. These findings are not unexpected given the central role for Pparα in regulating the hepatic genes necessary for fasting oxidative metabolism.

The loss of Pparα alone suppressed transcription of genes for fatty acid catabolism and ketogenesis, including *Ehhadh*, *Hmgcs2*, and *Fgf21* (**Fig. 3g**). Many of these genes displayed blunted expression in DKO livers as well (**Fig. 3h**). Curiously, several canonical Pparα target genes, such as *Plin2* and *Pdk4*, retained induction in DKO liver compared to control animals (**Fig. 3h**). Further analysis of the RNA-seq data revealed that Cpt2^L−/−^ and DKO animals shared 257 upregulated transcripts, respectively representing 42% and 40% of induced genes for those genotypes (**Fig. 3i**). This suggested that there may be a transcriptional program triggered by the shared metabolic perturbations presented by loss of hepatic β-oxidation. Further, 49% of induced transcripts in DKO are responsive to WY-14643 (**Fig. 3i**) (Naiman et al., 2019). We therefore hypothesized that a subset of genes, which are typically thought to rely upon Pparα for their induction, have in addition a secondary regulatory mechanism. The limited transcription of target genes that do not require Pparα is likely prompted by excess lipid burden via an independent mechanism driven by Err or Hnf4α, for example.

### Impaired hepatic fatty acid oxidation prompts distinct modes of gene regulation

We subsequently sought further resolution into the role for Pparα in regulating hepatic transcription in response to impaired hepatic fatty acid oxidation, including the possibility of Pparα-independent regulatory mechanisms. Unbiased k-means clustering was used to identify unique patterns of gene transcription across all differentially expressed transcripts (**Fig. S4a, Table S5**). Of the five clusters identified, two in particular (1 & 3) were associated with the Cpt2^L−/−^ gene induction signature (**Fig. 4a, 4d**). Notably, these two groups of transcripts display significantly divergent modes of Pparα-sensitive transcription.

**Fig 4.**
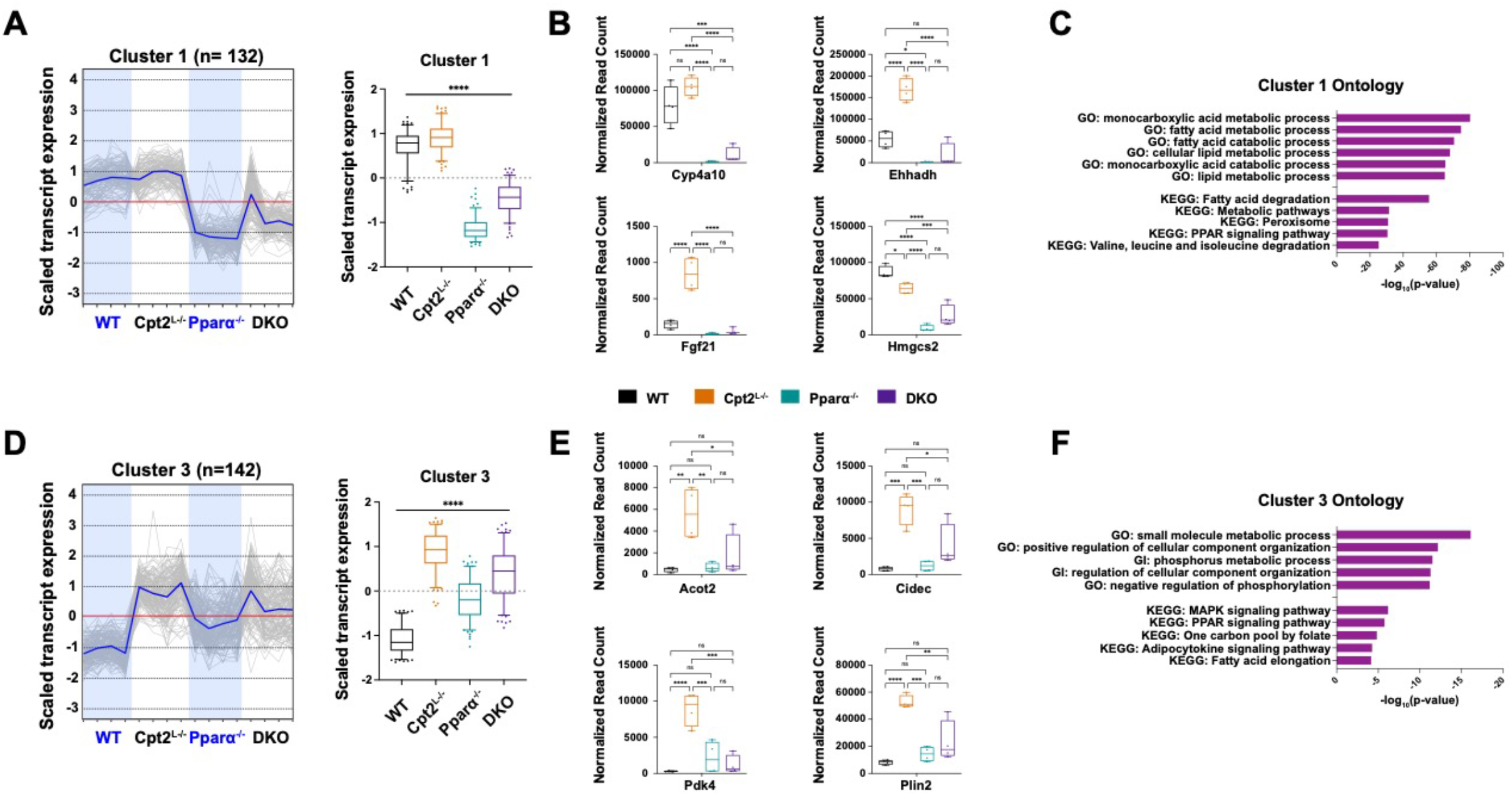
Unbiased clustering reveals patterns for Pparα-dependent and Pparα-independent transcription of target genes. A. *(left)* Tracing diagram of cluster 1 genes (Pparα-dependent) from fasted RNA-seq transcript levels. Blue line indicates mean expression. *(right)* Bar graph quantifying scaled transcript values. Presented as Z-score scaled transcript levels. B. Normalized transcript read counts for select cluster 1 genes. C. Gene ontology for cluster 1 genes ranked by significance. D. *(left)* Tracing diagram of cluster 3 genes (Pparα-independent) from fasted RNA-seq transcript levels. Blue line indicates mean expression. *(right)* Bar graph quantifying scaled transcript values. Presented as Z-score scaled transcript levels. E. Normalized transcript read counts for select cluster 3 genes. F. Gene ontology for cluster 3 genes ranked by significance. One-way ANOVA followed by Tukey’s post-hoc test was performed as appropriate. *p < 0.05; **p < 0.01; ***p < 0.001; ****p < 0.0001; ns, not significant. Bar graphs represent 2%-98% percentile.

Cluster 1 genes were wholly dependent upon Pparα for their transcription, as indicated by their repressed expression in Pparα^−/−^ and DKO fasted liver (**Fig. 4a**). Cluster 1 genes included multiple canonical Pparα targets, including *Cyp4a10*, *Ehhadh*, and *Hmgcs2* (**Fig. 4b, S3b**). These genes are respectively implicated in Pparα regulation of ω-oxidation, peroxisomal oxidation, and ketogenesis (Kersten et al., 2010). Indeed, gene ontology and KEGG pathway analysis on all Cluster 1 genes returned terms related to fatty acid metabolism and Pparα signaling (**Fig. 4c**). These data provide further evidence that impaired hepatic fatty acid oxidation activates Pparα-dependent transcription upon fasting.

Cluster 3 genes were likewise induced in Cpt2^L−/−^ mice in a Pparα-dependent manner. However, this group differs from the previous cluster in that both Pparα-null backgrounds retained baseline expression levels (**Fig. 4d**). Cluster 3 genes thus sustained transcriptional activity in the absence of Pparα, yet still required the transcription factor for induction in response to metabolic perturbations such as a buildup of Pparα ligands. This group included canonical Pparα target genes *Acot2* and *Pdk4*, whose expression in Pparα-null animals displayed no significant variation from control animals (**Fig. 4e**). KEGG pathway analysis returned Ppar Signaling Pathway as a top result (**Fig. 4f**). Curiously, Cluster 3 contained multiple genes associated with lipid droplets, including four members of the perilipin family (*Plin1-4*) and *Cidec* (Greenberg et al., 2011). *Plin2*, *Plin4*, and *Cidec* are known Pparα targets (Francque et al., 2015; Kersten et al., 2010). This suggests that modules of the Pparα transcription program involve building upon pre-established Pparα-independent gene expression patterns to respond to specific hepatocyte metabolic states.

### Distinct Pparα target genes retain promotor accessibility despite loss of Pparα

We next asked how these unique modes of Pparα target gene transcription were reflected at the level of chromatin architecture by assessing ATAC-seq from all four genotypes. Loss of Pparα drastically diminishes chromatin accessibility (**Fig. S5a**). Chromatin landscape remodeling for the Pparα-null mice was expected given the transcription factor’s crucial role in coordinating fasting metabolism, yet the severe ATAC-seq signal depletion in gene promoter regions was particularly striking (**Fig. 5a**). PCA emphasized this disparity, showing that Pparα and DKO animals clustered uniquely, while WT and Cpt2^L−/−^ livers overlapped substantially (**Fig. S5b**). We further observed there are appreciably fewer changes in discrete peak intensities between Pparα^−/−^ and DKO animals compared to Cpt2^L−/−^ and DKO animals (**Fig. 5b**). Together these data indicate that Pparα is required for proper upkeep of fasting chromatin dynamics.

**Fig 5.**
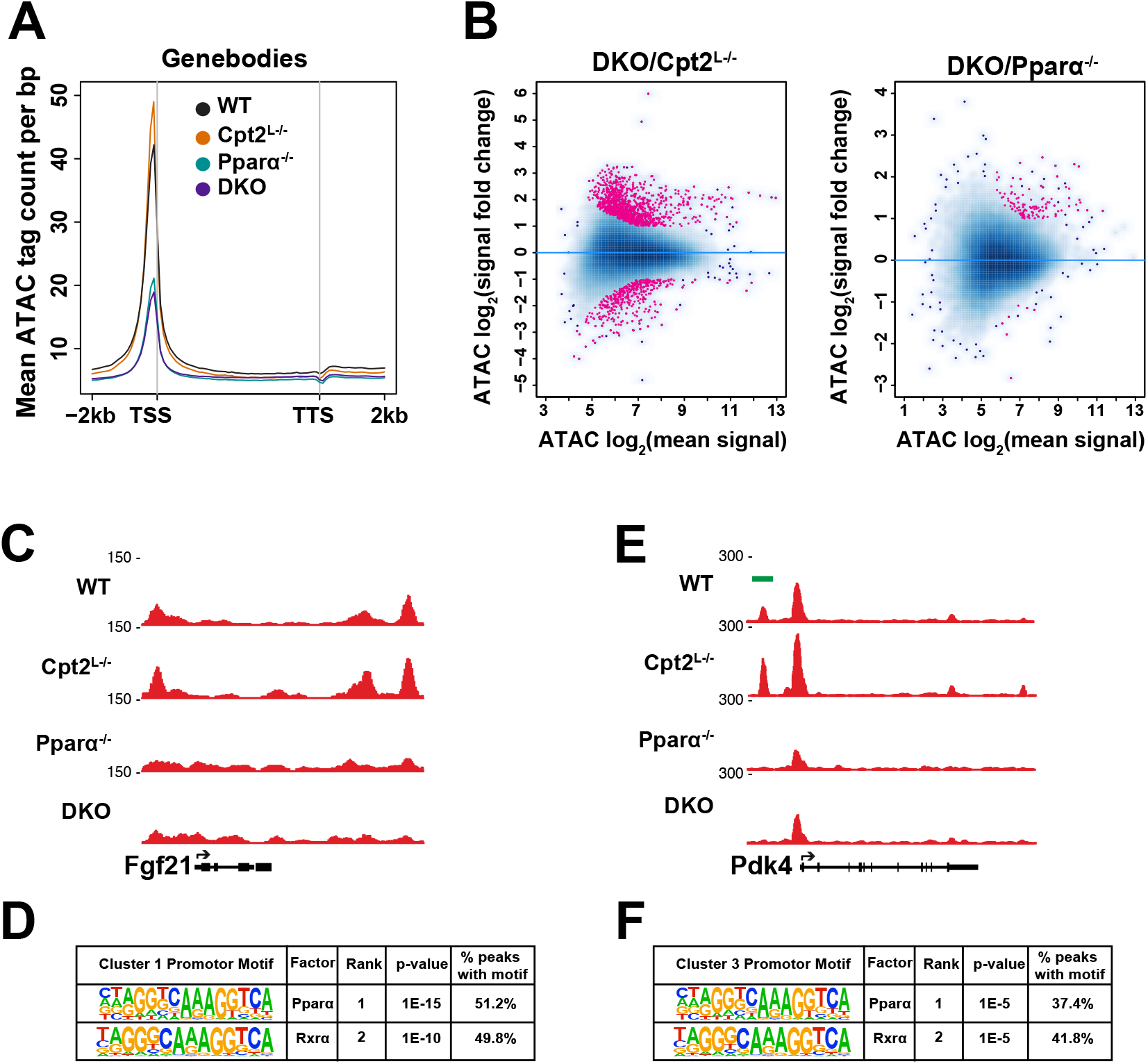
Loss of Pparα remodels genome-wide accessibility. A. Aggregation plot depicting liver ATAC-seq mean tag density for all gene bodies with ±2kb flanking regions in fasted WT, Cpt2^L−/−^, Pparα^−/−^, and DKO liver (n=2). TSS = transcription start site, TTS = transcription termination site. B. MA plot depicting ATAC-seq differential binding analysis results between DKO/Cpt2^L−/−^ and DKO/Pparα^−/−^ animals. The x-axis gives the mean signal and the y-axis gives the signal fold change for a given peak. Points in pink are significant (fold change ≥ |2|, FDR < 0.05). C. ATAC-seq genome browser tracks for cluster 1 gene *Fgf21*. D. HOMER enrichment analysis for known motifs in ATAC-seq promoter peaks for cluster 1 genes. E. ATAC-seq genome browser tracks for cluster 3 gene *Pdk4*. Pparα-sensitive peak interval is indicated by green bar. F. HOMER enrichment analysis for known motifs in ATAC-seq promoter peaks for cluster 3 genes.

Cluster 1 genes, those dependent upon Pparα for their expression, exhibited a common pattern of promotor chromatin accessibility that is represented by *Fgf21* (**Fig. 5c**). ATAC-seq genome browser tracks indicated a higher promoter peak signal in the Cpt2^L−/−^ livers compared to WT; accessibility was lost under a Pparα-null background. Indeed, motif analysis on ATAC-seq promotor peaks for this cluster returned Pparα and its heterodimer partner Rxrα as the top two hits (**Fig 5d**). Suppressed chromatin accessibility at loci enriched for Pparα binding motifs indicates the need for Pparα for proper promoter dynamics.

Promoter accessibility for Cluster 3 genes, those with Pparα-independent baseline expression, likewise shared a common promoter chromatin architecture as illustrated by *Pdk4* (**Fig. 5e**). The Cpt2^L−/−^ induction pattern was once again evident in the ATAC-seq signal. However, in contrast to Cluster 1 genes, this group retained a degree of promotor accessibility in Pparα^−/−^ and DKO mice. Interestingly, 13% of Cluster 3 genes contained an interval of open promotor chromatin that was lost in both Pparα-null animals, including at the promoters for Pparα targets *Acot2*, *Fabp3*, *Pdk4*, and *Plin2* (**Fig. 5e** green bar). These likely indicate binding sites for Pparα directly or for another DNA regulatory protein that depends on Pparα for an aspect of its activity. Cluster 3 ATAC-seq promoter peaks are enriched for Pparα and Rxrα binding motifs, albeit at a lower rate than for Cluster 1 genes (**Fig. 5f**). These data together indicate that Cluster 3 promoters do not require Pparα to maintain accessibility during a fast, though Pparα can interact with these loci to augment target gene transcription in response to a metabolic perturbation.

### Pparα-dependent and independent promotor acetylation

Differential promotor accessibility prompted questions as to how the loss of Pparα affected promotor dynamics *vis a vis* histone acetylation. We therefore performed ChIP-seq for H3K27ac, a histone mark generally found at active chromatin (Andersson and Sandelin, 2020). H3K27ac signal was comparable between genotypes, however Cpt2^L−/−^ and DKO livers displayed higher H3K27ac signal near transcription start sites compared to the WT and Pparα^−/−^ animals (**Fig. 6a, S6a**). PCA showed that WT and Cpt2^L−/−^ H3K27ac distributions were distinct from one another, while Pparα^−/−^ and DKO animals clustered together (**Fig. 6b**). This is the inverse of the ATAC-seq PCA, in which WT and Cpt2^L−/−^ animals were grouped and Pparα^−/−^ and DKO animals were separated (**Fig. S5b**). These data indicate that the metabolic perturbations in Cpt2^L−/−^ and DKO livers stimulate a common configuration of H3K27ac occupancy regardless of overall chromatin accessibility. This further demonstrates lipid-driven selective histone acetylation despite absence of mitochondrial β-oxidation. Pparα is not required for this H3K27ac deposition, however Pparα is still needed to promote chromatin accessibility and ultimately transcription of lipid catabolic genes. Comparing the WT and Cpt2^L−/−^ ATAC-seq and H3K27ac ChIP-seq peaksets provides additional evidence that the Cpt2^L−/−^ epigenetic program builds upon the baseline fasting chromatin architecture to exaggerate an established response.

**Fig 6.**
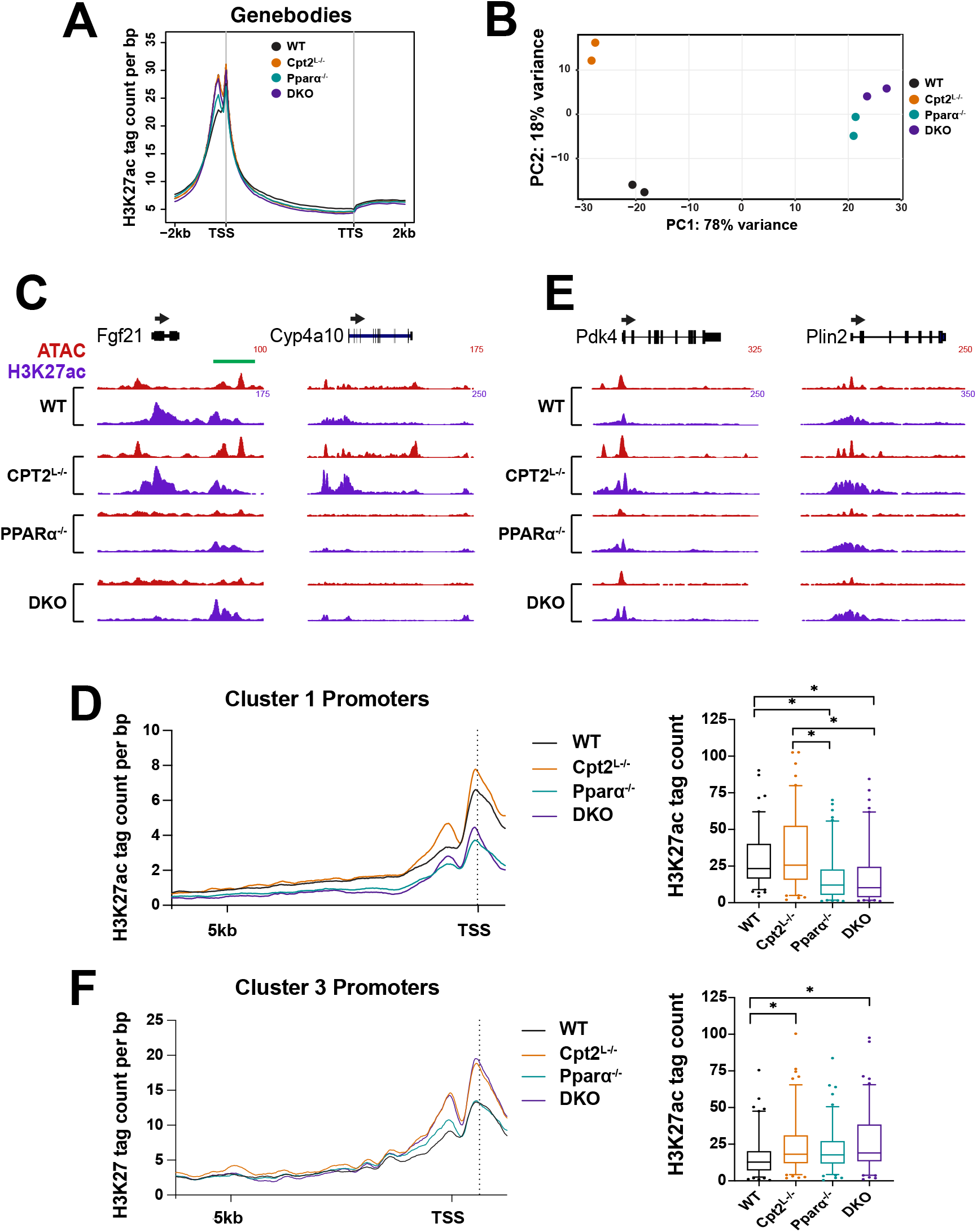
Differential H3K27ac occupancy patterns at Pparα target genes. A. Aggregation plot depicting liver H3K27ac ChIP-seq mean tag density for all gene bodies with ±2kb flanking regions in fasted WT, Cpt2^L−/−^, Pparα^−/−^, and DKO liver (n=2). TSS = transcription start site, TTS = transcription termination site. B. PCA of H3K27ac ChIP-seq data from fasted WT, Cpt2^L−/−^, Pparα^−/−^, and DKO liver. C. Genome browser track for cluster 1 genes *Fgf21* and *Cyp4a10*. ATAC-seq is presented in red, H3K27ac ChIP-seq is presented in purple. Green bar indicates downstream regulatory region. D. (*left*) Aggregation plot for H3K27ac ChIP-seq mean tag density for cluster 1 gene promoters. (*right*) Bar graph quantifying mean values of H3K27ac ChIP-seq peaks found in cluster 1 gene promoters. E. Genome browser track for cluster 3 genes *Pdk4* and *Plin2*. ATAC-seq is presented in red, H3K27ac ChIP-seq is presented in purple. F. (*left*) Aggregation plot for H3K27ac ChIP-seq mean tag density for cluster 3 gene promoters. (*right*) Bar graph quantifying mean values of H3K27ac ChIP-seq peaks found in cluster 3 gene promoters. Significance determined by Kruskal-Wallis test with Dunn’s post hoc correction. *p < 0.0001; ns, not significant. Bar graphs represent 2%-98% percentile.

Clusters 1 and 3, together comprising the genes induced in Cpt2^L−/−^ liver, provided a useful contrast to understand variations in active chromatin acetylation and how they relate to Pparα. H3K27ac occupancy at Cluster 1 gene promotors paralleled ATAC-seq trends in that loss of Pparα was associated with significantly decreased promoter H3K27ac tag density, as indicated by *Fgf21* and *Cyp4a10* (**Fig. 6c**). There was no statistical difference in H3K27ac signal between WT and Cpt2^L−/−^ animals (**Fig. 6d**). At Cluster 3 gene promoters, in contrast to Cluster 1, Pparα loss was not associated with depleted promoter H3K27ac signal. H3K27ac occupancy was instead distinctly independent of Pparα, as indicated by its canonical target genes *Pdk4* and *Plin2* (**Fig 6e**). Moreover, the Cpt2^L−/−^ background is associated with significantly elevated H3K27ac promoter tag density compared to WT and Pparα^−/−^ animals (**Fig. 6f**). These observations continue to demonstrate that the lipid-associated changes in chromatin accessibility are linked to site-specific epigenetic modifications in gene promoters.

At Cluster 1 gene *Fgf21* we noticed an acetylation pattern indicative of a regulatory region downstream from the transcription termination site (**Fig. 6c**, green bar). The H3K27ac signal at this site was comparable between the four genotypes, yet accessibility was only observed in the WT and Cpt2^L−/−^ animals. Both ATAC-seq peaks contained Pparα binding motifs. This chromatin pattern parallels that observed at cluster 3 gene promoters. We speculate that at these loci Pparα is able to detect antecedent chromatin acetylation as part of its DNA binding mechanism, or that an epigenetic reader detects that mark and subsequently loads Pparα onto chromatin, thereby promoting a transcriptionally permissive state.

### Loss of Pparα selectively alters the hepatic fasting enhancer landscape

Fasting stimulates a cascade of transcription factor loading and binding onto hepatic enhancers (Goldstein et al., 2017). Enhancers can be categorized as either poised or active. The former are indicated by the histone modification H3K4me1, and the latter are defined by co-occupancy of H3K4me1 and H3K27ac (Creyghton et al., 2010). We decided to use the Cpt2^L−/−^ model as a handle by which to investigate lipid-responsive enhancer elements, and the extent to which Pparα regulates both their poising and activation. A total of 8998 active enhancers were detected across all genotypes (**Fig. 7a, Table S6**). Strikingly, Pparα-null animals had diminished chromatin accessibility over active enhancer elements (**Fig. S7a**). Loss of either Cpt2 or Pparα alone did not impact H3K4me1 density at active enhancers, but their combined deletion in DKO liver reduced global enhancer priming (**Fig. 7a**). Enhancer H3K27ac signal did not differ between Pparα^−/−^ and WT animals. However, loss of hepatic Cpt2 was associated with overall decreased H3K27ac occupancy at active enhancers. This was an unanticipated contrast to increased acetylation at gene promoters under the Cpt2^L−/−^ background (**Fig. 1g, 6a**). Overall these data imply that while loss of Pparα alone does not negatively impact the enhancer epigenetic landscape during the latter stages of a fast, global enhancer accessibility is significantly hindered without the transcription factor.

**Fig 7.**
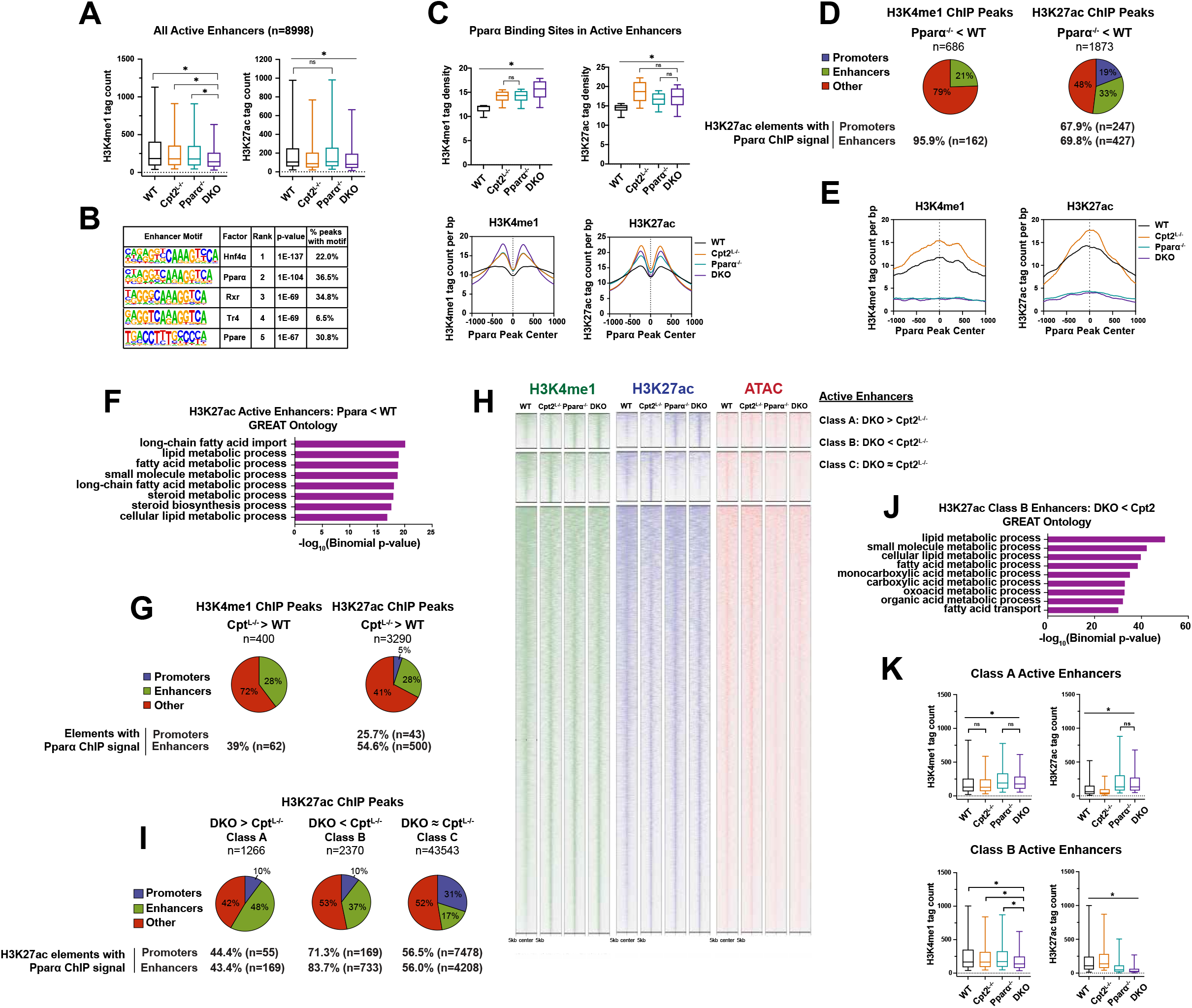
Lipid-sensitive enhancer elements are primed for Pparα. A. Bar graph depicting H3K4me1 and H3K27ac tag counts at active enhancer elements. B. HOMER motif analysis for all active enhancers as defined by H3K4me1 and H3K27ac co-occupancy. C. (*top*) Bar graphs depicting H3K4me1 and H3K27ac mean tag density proximal to Pparα ChIP-seq binding sites in all active enhancers (±500bp from Pparα ChIP peak center). (*bottom*) Aggregation plot for H3K4me1 and H3K27ac tag density within 1kb flanking regions of Pparα ChIP peak center. (Pparα ChIP-seq: GSE118788, Sommars et al., 2019) D. (*top*) Pie chart showing genomic distribution (promoters, enhancers, other) of H3K4me1 and H3K27ac differential peak analysis at enhancers with repressed H3K27ac signal in Pparα liver compared to WT. Promoters were not considered for H3K4me1 ChIP-seq. (*bottom*) Percent overlap between peak genomic annotation and Pparα ChIP-seq coordinates. E. Aggregation plots for H3K4me1 and H3K27ac tag density within 1kb flanking regions of Pparα ChIP peak center at enhancers with repressed H3K27ac signal in Pparα liver. F. H3K27ac GREAT ontology for enhancers with repressed H3K27ac signal in Pparα liver. G. (*top*) Pie chart showing genomic distribution of H3K4me1 and H3K27ac differential peak analysis for enhancers with increased H3K27ac signal in Cpt2^L−/−^ liver compared to WT. (*bottom*) Percent overlap between peak genomic annotation and Pparα ChIP-seq coordinates. H. Heatmap of normalized H3K27ac ChIP-seq (purple), H3K4me1 ChIP-seq (green), and ATAC-seq (red) signal at active enhancers classified according to DKO/Cpt2^L−/−^ differential peak analysis. Coverage is calculated from peak center with ±5kb flanking regions. I. (*top*) Pie chart showing genomic distribution of H3K27ac differential peak analysis for enhancers with increased, repressed, and unchanged H3K27ac signal in DKO liver compared to Cpt2^L−/−^. *(bottom*) Percent overlap between peak genomic annotation and Pparα ChIP-seq coordinates. J. H3K27ac GREAT ontology for Class B enhancers (DKO < Cpt2^L−/−^). K. Bar graphs quantifying normalized H3K27ac and H3K4me1 ChIP-seq signal at Class A and Class B active enhancers Significance determined by Kruskal-Wallis test with Dunn’s post hoc correction. *p < 0.0001; ns, not significant. Bar graphs represent 2%-98% percentile.

Active enhancers were highly enriched for Hnf4α and Pparα binding motifs (**Fig. 7b**). Surprisingly, Pparα ChIP-seq sites located within enhancers showed elevated H3K4me1 and H3K27ac signals in all knockout genotypes (**Fig. 7c**) (Sommars et al., 2019). There was no difference in enhancer H3K4me1 signal between Cpt2^L−/−^ and Pparα^−/−^ liver, though loss of both genes in DKO animals caused significantly increased methylation near Pparα binding sites.

H3K27ac tag density flanking Pparα binding sites was increased in Cpt2^L−/−^ animals compared to Pparα^−/−^. Cpt2^L−/−^ animals also had increased ATAC-seq signal over Pparα enhancer sites compared to WT animals (**Fig. S7b**). As might be expected, chromatin accessibility over Pparα sites was decreased in Pparα-null livers, though this was despite increased enhancer priming and acetylation over WT baselines at those loci (**Fig. 7c, S7b**). This indicates that the active epigenetic state is maintained independent of Pparα at these enhancers, but Pparα is required to promote chromatin accessibility. Together these data show that perturbations to fatty acid catabolism are associated with amplified enhancer priming and acetylation specifically near Pparα binding sites even in the absence of the transcription factor. We propose this is an epigenetic mechanism for the hepatocyte to prime a genomic response to impaired lipid catabolism.

We next examined the specific changes to enhancer state at loci that are differentially regulated in Pparα^−/−^ liver compared to WT. Loss of Pparα was associated with targeted repression of H3K27ac signal, and to a lesser extent H3K4me1 signal (**Fig. 7d**). Only 169 enhancers had suppressed priming in Pparα^−/−^ animals in contrast to the 612 enhancers with suppressed H3K27ac signal in Pparα^−/−^ animals. As expected, these downregulated regions contained a high frequency of Pparα ChIP-seq binding sites. Curiously, 33% of H3K27ac peaks repressed in Pparα^−/−^ liver were located within enhancer elements, while only 19% of H3K27ac signal changes were found in gene promoters (**Fig. 7d**). In other words, loss of Pparα diminished chromatin acetylation at a greater number of enhancers than promoters. Enhancers with blunted acetylation in Pparα^−/−^ liver displayed highly enriched H3K4me1 and H3K27ac signals in Cpt2^L−/−^ animals (**Fig. 7e**). The Genomic Regions Enrichment of Annotations Tool (GREAT), which provides ontologies for enhancer function using a binomial test specific for long-range genomic regulatory domains, showed that enhancers repressed by loss of Pparα were associated with ontology terms for fatty acid metabolism (**Fig. 7f**) (McLean et al., 2010). These data are consistent with a model in which Cpt2 loss drives Pparα signaling, and that Pparα target gene transcription is associated with heightened activation of enhancers associated with lipid metabolism.

Following up on these observations, we next examined how a deficit in β-oxidation affected the hepatic enhancer landscape, and the extent to which Pparα is implicated in genomic maintenance under these metabolic conditions. Loss of Cpt2 had minimal effects on overall enhancer priming (**Fig. 7g**). In contrast, Cpt2^L−/−^ mice had 915 enhancer regions with increased H3K27ac signal. 55% of these regions contain Pparα binding sites. Indeed, Cpt2^L−/−^ animals had a prominent increase in both enhancer priming and acetylation over Pparα ChIP sites (**Fig. S7c**). At these Cpt2^L−/−^-sensitive loci we also observed increased H3K4me1 and H3K27ac ChIP signals over Pparα binding sites in Pparα-null animals compared to WT controls, indicating that enhancers with epigenetic sensitivity to impaired β-oxidation likewise respond to impaired Pparα signaling, perhaps designating a response to drive augmented transcription such as we see in the Cpt2^L−/−^ mice (**Fig. S7c**).

The DKO mice clarify the role for Pparα at these enhancer elements. These mice had a striking change to their active enhancer landscape when compared to their Cpt2^L−/−^counterparts. We initially noticed there were far fewer changes in enhancer priming compared to changes in enhancer acetylation (**Fig. 7i, S7d**). A total of 1487 differentially regulated enhancers were detected between the two genotypes, which were broken down into three groups according to differential H3K27ac ChIP signal (**Fig. 7h**). Class B enhancers (n=876) were defined by statistically significant H3K27ac depletion in DKO mice compared to Cpt2^L−/−^ (**Fig. 7h, 7k**). We consider these to be the enhancer elements at which the ligand-activated Pparα signaling program specifically impacts acetylation in response to lipid sensing. The high incidence of Pparα binding sites within these enhancers suggests that Pparα itself plays a direct role in local H3K27ac deposition (**Fig. 7i**). Cpt2^L−/−^ animals had increased H3K27ac signal at these regions compared to WT (**Fig. 7k**). We further noted that loss of Pparα on the Cpt2^L−/−^ background had an outsized effect on enhancer acetylation compared to promoter acetylation (**Fig. 7i**). Finally, GREAT analysis indicates that proximal genes for Class B enhancers were associated with multiple lipid gene ontology terms (**Fig. 7j**). Together these data demonstrate the substantial role for enhancer elements in the Pparα transcriptional response, and that lipid-amplified Pparα signaling is perhaps mediated more substantially by epigenetic changes to enhancer regions than at promoters.

Class A enhancers (n=611) were defined by a greater than two-fold increase in H3K27ac signal in DKO mice compared to Cpt2^L−/−^ liver. DKO Class A enhancer acetylation was similarly increased over WT animals (**Fig. 7k**). This augmented H3K27ac signal was shared by Pparα^−/−^ animals. Likewise, both Pparα^−/−^ and DKO animals displayed elevated H3K4me1 priming levels at Class A enhancers (**Fig. 7k**). This implies Pparα loss is associated with enhancer activation that may be linked to other metabolic processes. Class C enhancers (n=7511), which showed no change in acetylation state between Cpt2^L−/−^ and DKO animals, represent 83% of the active enhancer landscape (**Fig. S7e**), emphasizing that the genomic perturbations presented by loss of Cpt2 or Pparα result in specific, targeted changes to enhancer epigenetic state.

Altogether these data demonstrate that metabolic perturbations such as inhibiting hepatic fatty acid oxidation or loss of Pparα result in higher levels of enhancer priming and acetylation proximal to Pparα sites that are largely associated with lipid metabolism. Pparα DNA binding is proceeded by enhancer priming and acetylation. This is clearly illustrated at Pparα-bound enhancers found near *Fgf21* and *Pdk4* (**Fig. S7f**). The acetylation may even be the signal for Pparα to engage these regulatory elements. Furthermore, while changes to enhancer epigenetic state are more localized near Pparα binding sites, the transcription factor is so crucial to maintaining permissive chromatin that loss of Pparα repressed chromatin accessibility across the global enhancer network. We also surmise that changes to the enhancer landscape comprise a significant component of ligand-activated Pparα signaling, with quantitatively more Pparα-associated changes in acetylation occurring at enhancers compared to promoters. Overall, these data show the requirement for Pparα activation in maintaining transcriptionally permissive hepatic genomic architecture, particularly when the hepatocyte must sense and respond to elevated lipid.

## DISCUSSION

A foundational question in liver metabolism is how the hepatocyte senses and responds to fluctuations in lipid availability. This ultimately requires tight coordination between the hepatocyte’s metabolic state, genome architecture, and transcriptional output. Fasting Cpt2^L−/−^ mice prompts accumulation of fatty acids, including putative endogenous Pparα ligands (Lee et al., 2016; Selen et al., 2021). Using this mouse genetic model and a combination of next-generation sequencing platforms we clarify the role for hepatic lipid content in metabolic gene transcription, and in particular how Pparα, activated by natural ligands, maintains the transcriptional architecture necessary for target gene expression (**Fig. 8**).

**Fig 8.**
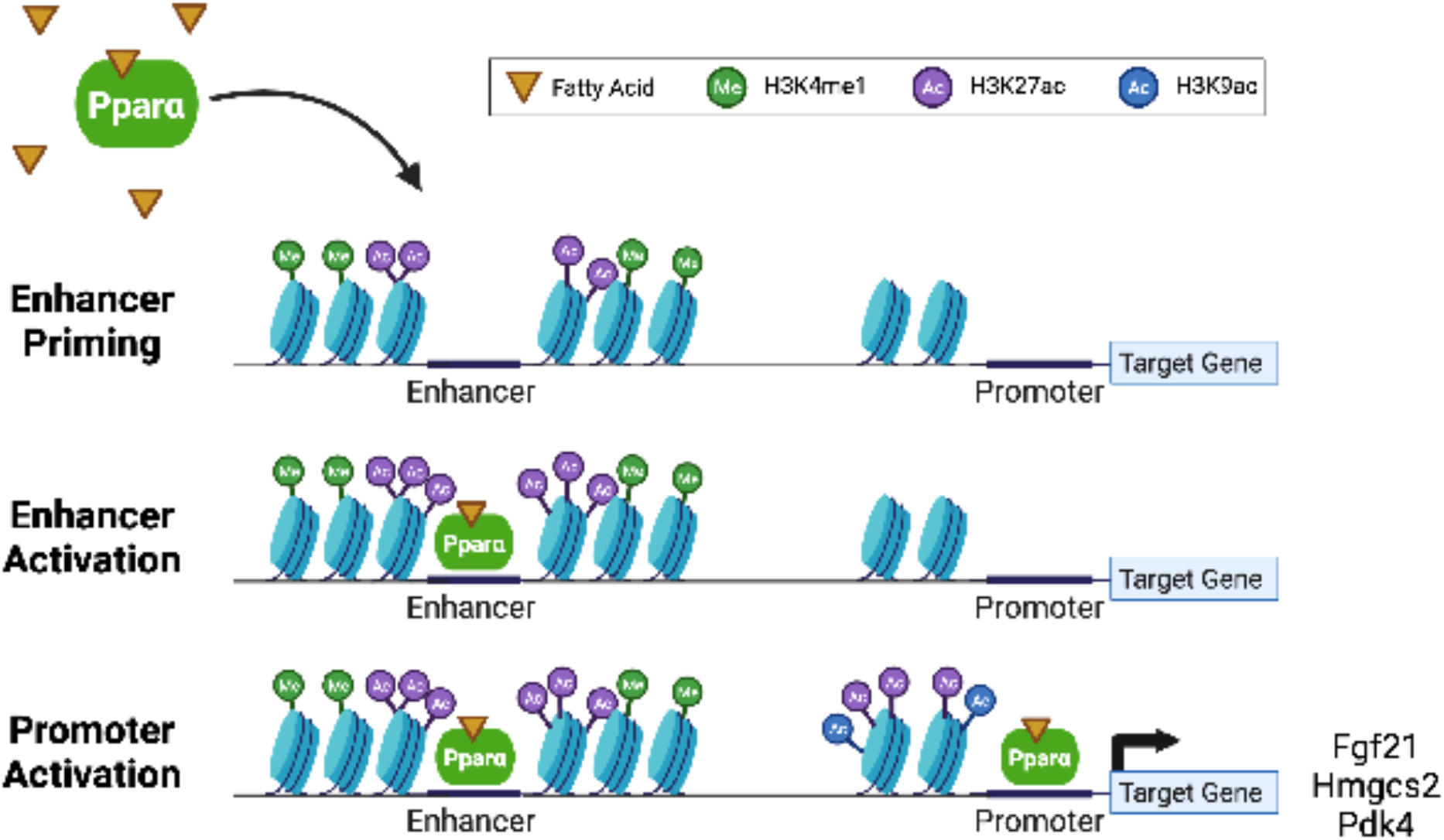
Pparα is required for lipid-responsive enhancer and promoter activation. Deficit in mitochondrial β-oxidation and/or excess fatty acids prompts H3K4me1 and H3K27ac deposition at lipid-responsive enhancer elements. This signals ligand-activated Pparα recruitment, and subsequent enhancer accessibility and activity. Pparα then binds to target gene promoters to drive chromatin acetylation, thereby inducing transcription of fatty acid catabolic genes to alleviate heightened lipid burden.

Our system benefits from use of *in vivo* metabolic perturbations to assess the role for excess fatty acids in driving transcription. It has been stipulated that fatty acid oxidation promotes transcription of metabolic genes via bulk increase to histone acetylation by lipid-derived acetyl-CoA (McDonnell et al., 2016). Our data instead reveal that changes in hepatic lipid levels can induce metabolic gene transcription even when mitochondrial β-oxidation, and subsequent acetyl-CoA production, is inhibited. Indeed we detected no changes in bulk histone acetylation in Cpt2^L−/−^ liver. We instead observed that promoter acetylation patterns for many lipid catabolic genes are Pparα-dependent. Enhancers displayed Pparα-independent acetylation proximal to Pparα binding sites, but enhancer chromatin remained inaccessible in the absence of the transcription factor. This indicates that fatty acid-induced transcription is stimulated not by epigenetic changes, but instead by ligand-induced Pparα activation and the associated chromatin remodeling. Studies in mice given a high fat diet (HFD) provide supplemental evidence for our observations on fatty acid-driven transcription. In parallel with our results, HFD-fed mice show increased hepatic lipid content as well as Pparα-dependent hepatic gene expression (Patsouris et al., 2006). HFD-fed mice likewise exhibit suppressed liver acetyl-CoA content but no difference in bulk histone acetylation compared to chow-fed controls (Carrer et al., 2017).

A common method to stimulate Pparα and define target genes is to use a pharmacological agonist (Brocker et al., 2020; Janssen et al., 2015; Lee et al., 1995; Rakhshandehroo et al., 2007). Here we instead use fasting-induced buildup of endogenous ligand to activate Pparα. We observed two distinct patterns of target gene transcription. In the first group (Cluster 1), target genes require Pparα for promoter accessibility and acetylation under conditions of high lipid availability. In the second group (Cluster 3), target genes do not require Pparα for promoter acetylation. Promoter accessibility for the latter cluster is maintained in Pparα^−/−^ animals, though to a lesser extent than measured in WT controls. We therefore hypothesize that a secondary mechanism maintains transcriptionally permissive H3K27ac occupancy at these loci independent of Pparα, thereby promoting baseline gene expression. The additional requirement of Pparα for gene induction above baseline is reflected in both the RNA-seq and ATAC-seq datasets.

Cluster 3 gene promoters are enriched for the Hnf4α biding motif, suggesting the nuclear hormone receptor may be involved in Pparα-independent promoter acetylation. It should be noted that Hnf4α can bind directly to promoter sequences of *Fgf21*, *Hmgcs2*, and *Cpt1* to obstruct Pparα binding and gene transcription (Martinez-Jimenez et al., 2010). However, these three genes were assigned to Cluster 1, leaving open the possibility of Hnf4α involvement in Cluster 3 promoter activation. Hnf4α is highly expressed in the three knockout genotypes, providing additional evidence for this suggestion. We similarly observe high incidence of Hnf4α motifs at active enhancers, in line with previous work demonstrating that Hnf4α maintains H3K27ac occupancy at active enhancers in mouse liver (Thakur et al., 2019).

Previous *in vivo* work has advanced a role for Pparα localization to hepatic enhancers. In a model of diet induced obesity, Pparα-null mice had suppressed circadian transcription for eRNA proximal to Pparα binding motifs (Guan et al., 2018). In another study, hepatic fed state Pparα ChIP-seq binding sites were observed to overlap with fasting-induced enhancers (Goldstein et al., 2017). Our research builds upon those observations by demonstrating that Pparα-sensitive enhancer elements in fasting liver are associated with lipid signaling. Elevated enhancer accessibility in Cpt2^L−/−^ animals, including particularly high ATAC-seq signal over Pparα binding sites, further indicates that lipids drive Pparα engagement with active chromatin. Moreover, depleted chromatin accessibility over all active enhancers in Pparα-null animals indicates the requirement of Pparα to properly maintain the hepatic fasting enhancer landscape.

It’s notable that Pparα loss does not significantly impact global enhancer epigenetic state. Indeed, enhancer priming proximal to Pparα binding sites is independent of Pparα itself. Increased H3K4me1 signal in DKO animals compared to other genotypes likely indicates a compensatory signal to engage lipid-sensitive enhancers in response to the hepatocyte’s need for oxidative clearance of excess fatty acid. Enhancer H3K27ac signal adjacent to Pparα binding sites is likewise globally independent of Pparα. Both Cpt2^L−/−^ and DKO animals display the highest levels of enhancer acetylation near Pparα binding sites, and yet these two genotypes also display the lowest global enhancer H3K27ac signal. In other words, the Cpt2^L−/−^ background drives enhancer acetylation specifically in proximity to Pparα binding sites, providing evidence for a mechanism in which high lipid levels prompt H3K4me1 and H3K27ac deposition to prime Pparα-sensitive enhancers, and credibly acting as a signal for Pparα DNA binding.

While overall enhancer maintenance near Pparα binding sites is independent of Pparα, insight into the Pparα enhancer program is gained by examining differential comparisons between genotypes. One striking observation was that Pparα-associated H3K27ac signal changes occurred more frequently in enhancer elements compared to promoters. Cpt2^L−/−^-associated lipid prompts an H3K27ac ChIP-seq signal increase at 5.5x more enhancers than promoters. Similarly, loss of Pparα in Class B enhancers (DKO < Cpt2^L−/−^) is associated with H3K27ac signal depletion at 3.7x more enhancers than promoters. Finally, Pparα^−/−^ animals show H3K27ac depletion at 1.7x more enhancers than promoters. This notable degree of signal change at enhancer regions, combined with the high incidence of Pparα binding sites and association with GREAT lipid metabolism ontology terms, suggests that Pparα effects more transcriptionally influential changes to chromatin architecture at enhancers over promoters. We believe this indicates that the mechanism for adjusting Pparα target gene transcription levels is more closely linked to enhancer remodeling than to promoter dynamics.

These expanded insights into Pparα gene regulation provide evidence for discrete modes by which the liver signals its transcriptional needs in the face of metabolic perturbations linked to fluctuations in lipid availability. We demonstrate distinct types of Pparα signaling based on patterns of chromatin state and transcriptional output, show the role for Pparα in hepatic enhancer maintenance, and further suggest secondary mechanisms which aid and potentially even modulate Pparα signaling. These data prompt questions as to what role Pparα may play in regulating the transcriptional landscape associated with lipid sensing in other tissues, such as the kidney and adipose. This study thus uncovers new avenues of investigation into lipid sensing at the levels of locus-specificity, tissue-specificity, and broader metabolic physiology.

## MATERIALS & METHODS

### Animals

All procedures were performed in accordance with the NIH’s Guide for the Care and Use of Laboratory Animals and under the approval of the Johns Hopkins School of Medicine Animal Care and Use Committee.

Cpt2^fl/fl^ and Albumin-Cre;Cpt2^fl/fl^ (Cpt2^L−/−^) mice were previously described (Lee et al., 2015, 2016). Cpt2^fl/fl^;Pparα^−/−^ and Cpt2^L−/−^;Pparα^−/−^ animals were generated by initially crossing Cpt2^fl/fl^ or Cpt2^L−/−^ animals with Pparα^−/−^ mice (Jackson Laboratories; stock no. 008154). Mice for experiments were bred from either Cpt2^fl/fl^ crossed to Cpt2^L−/−^ animals or Cpt2^fl/fl^;Pparα^−/−^ crossed to Cpt2^L−/−^;Pparα^−/−^ animals. All mice were housed in a facility with ventilated racks on a 14h light/10h dark cycle with *ad libitum* access to a standard rodent chow (2018SX Teklad Global, 18% protein). For fasting experiments, 9 week old male mice were deprived of food for 24 hours (3p.m.-3 p.m). Tissues and serum were collected and flash-frozen at time of harvest.

### Immunoblotting

Flash-frozen liver tissue was homogenized in RIPA buffer (50 mM Tris-HCl at pH 7.4, 150 mM NaCl, 1 mM EDTA, 1% Triton X-100, and 0.25% deoxycholate) with PhosSTOP phosphatase Inhibitor (Roche) and protease inhibitor cocktail (Roche). Homogenates were rotated for 30 minutes at 4°C and then centrifuged at 18,000g for 15 minutes. Total protein concentrations were quantified by BCA assay (Thermo Fisher Scientific).

Protein lysates (30ug input) were separated by Tris-glycine SDS-PAGE (10-12% polyacrylamide gels), followed by a transfer to PVDF membranes (Immobilon). Membranes were blocked with 5% nonfat milk in TBST for 1 hour and incubated overnight at 4°C with primary antibodies at 1:1000 in 3% BSA in TBST: Acetylated-Lysine, 9441, Cell Signaling Technologies; Histone H3, 4499, Cell Signaling Technology, Acetyl-Histone H3 Lys9, 9649, Cell Signaling Technologies; Acetyl-Histone H3 Lys14, 7627, Cell Signaling Technologies; HSC70, 7298, Santa Cruz Biotechnology. HRP-conjugated anti-rabbit (Cytiva, NA934V) or fluorescence-based (Cy3-conjugated anti-mouse or Cy5-conjugated anti-rabbit; Invitrogen, Thermo Fisher Scientific) secondaries were used at 1:1000. Blots were imaged using the Amersham Prime enhanced chemiluminescent substrate (Cytiva) or epifluorescence on an Alpha Innotech MultiImage III instrument.

Histones were acid precipitated as previously described using an overnight 0.4N H_2_SO_4_ incubation and 1 hour TCA precipitation (Shechter et al., 2007). Histones were visualized per above using 15ug protein input on an 8% polyacrylamide gel.

### Mass Spectrometry & Proteomic Analysis

Flash-liver tissue was submitted to the Johns Hopkins Center for Proteomics Core for total proteome and acetyl-proteome analysis. Samples were prepared in lysis buffer with 8M urea and 50mM tetraethyl ammoniumbicarbonate. Lysates were treated with LysC and trypsin, then labeled with 10-plex TMT. Peptide fractionation was done using basic pH reverse phase liquid chromatography, resulting in 24 fractions. Fractions were analyzed using an Orbitrap Fusion Lumos (Thermo Scientific) on an Easy nLC 1200 (Thermo Scientific) using MS1 resolution = 120,000 and MS2 resolution = 30,000, HCD fragmentation method, and MS2 collision energy = 35. Each fraction received a two hour run time.

Total proteome analysis was done in Perseus (version 1.6.0.0) (Cox and Mann, 2012). p-value was calculated by Student’s t-test. q-value was calculated by significance analysis of microarrays (SAM) and permutation based false discovery rate (FDR), with SAM S0 value = 0.1 (Tusher et al., 2001).

Acetyl-proteome was analyzed with Proteome Discoverer 2.1 (Thermo Scientific) using RefSeq version 78. The following parameters were used: cleavage enzyme was set as trypsin, no more than five missed cleavages were allowed, fixed modification was carbamidomethyl on cysteine residue, dynamic modifications were acetyl group on protein N-terminus or oxidation on methionine and acetyl-lysine, and MS2 level quantification. A 1% FDR was applied for both peptide and protein levels. Acetyl-proteome data were normalized to median of total proteome to remove systemic deviation.

The COMPARTMENT dataset was used to assign localization for peptides for all cell compartments except mitochondria (Binder et al., 2014). Only COMPARTMENTS annotations with a minimum confidence score of 3 were used. Mitochondrial peptides were assigned using the MitoCarta 2.0 (Calvo et al., 2016).

### Tissue acetyl-CoA

Flash-frozen liver tissue was submitted to the Mouse Metabolic Phenotyping Center at Case Western Reserve University for acetyl-CoA measurements via liquid chromatography-mass spectrometry.

### Serum metabolites

Blood glucose levels in fasted mice were measured at time of harvest using a Nova Max Plus glucometer.

Untargeted metabolomics on serum was performed by Metabolon Inc (n=6). For analysis raw area counts for each biochemical species were rescaled to set the median equal to 1. Heatmap and PCA were generated by MetaboAnalyst (Chong et al., 2019). Differentially regulated metabolites for heatmap were determined using a 1-way ANOVA with Fisher’s LSD.

### RNA-sequencing library preparation

Total RNA was isolated from flash-frozen liver tissue was using TRIzol reagent (Invitrogen, Thermo Fisher Scientific), followed by addition purification using RNeasy Mini Kit (QIAGEN), per manufacturer recommendations. RNA quality was assessed by Nanodrop.

RNA was then submitted to Novogene Corporation Inc (China & Davis, CA, USA) for library construction and sequencing. Four biological replicates were used for each genotype. Reads were aligned to mouse reference genome (mm10) using STAR software (Dobin et al., 2013). Investigators performed analyses downstream of read count acquisition.

### RNA-seq Analysis

Differential expression was performed on raw read counts in R with DESeq2 (v3.12) using the Wald test with betaPrior=FALSE and lfcShrink type=”apeglm” (Love et al., 2014; Zhu et al., 2019). Differential expression cutoff was fold change ≥ |2|, padj < 0.05. Normalized read counts were visualized in GraphPad Prism. PCA plot was generated using pcaExlorer package (Marini and Binder, 2019). WY14643-responsive genes were determined from GSE140063 using the same differential expression pipeline (Naiman et al., 2019).

K-means clustering of differentially expressed transcripts was performed in R on rlog transformed count values from DESeq2 output. Optimal cluster number (n=5) was determined by gap statistic (factoextra package). K-means clustering was performed and visualized in the pheatmap package with Z-score row scaling 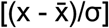. Tracing diagrams were generated using the ggplot2 package (Wickham, 2016). Gene ontology and KEGG pathway enrichment was obtained using the DAVID web application (Huang et al., 2007).

### ATAC-seq and ChIP-seq library preparation

ATAC-seq and ChIP-seq were performed using Active Motif sequencing services (Carlsbad, CA). Flash-frozen liver tissue was submitted to Active Motif, with two biological replicates per genotype for H3K4me1 and H3K27ac ChIP-seq, and ATAC-seq. One biological replicate was submitted for H3K9ac ChIP-seq. Library construction was performed according to company protocol. The following antibodies were used for ChIP-seq: H3K4me1 (Active Motif #39297), H3K9ac (Active Motif #39917), H3K27ac (Active Motif #39133).

### ATAC-Seq & ChIP-seq Analysis

Sequence acquisition, mapping, raw BAM file generation, and peak calling was performed by Active Motif. For ATAC-seq, paired-end 42nt sequencing reads generated by Illumina NexSeq 500 were mapped to mm10 reference genome using the BWA algorithm with default settings (Li and Durbin, 2009). For ChIP-seq, single-end 75nt sequencing reads generated by Illumina NexSeq 500 were mapped to mm10 reference genome using the BWA algorithm with “bwaln/samse” default settings. For normalization, tag number of all samples within a comparison group was reduced by random down-sampling to the number of tags present in the smallest sample. Peak calling was performed using MACS2 with default cutoff values (Zhang et al., 2008); ATAC-seq paired reads were treated as independent events. ENCODE blacklist regions were removed (Amemiya et al., 2019). bigWig files for visualization on UCSC Genome Browser were created using a 32nt bin with a 200bp *in silico* tag extension (Kent et al., 2002). Investigators performed downstream analysis.

Raw BAM files were processed using samtools and bamtools to remove reads with mapq score < 25, unmapped reads, mitochondrial reads, and reads with more than 2 mismatches (Barnett et al., 2011; Li et al., 2009). Aggregation plots and motif analysis were generated using Hypergeometric Optimization of Motif EnRichment (HOMER) (Heinz et al., 2010). Motif analysis was calculated using ‘findMotifsGenome.pl’ with-size 500 on BED file input with that assumption that a broad scanning window is preferred for histone mark motif analysis. knownResults motif output are reported in this study. Tag directories for histograms were generated from BAM files using ‘makeTagDirectory.pl’ with -fragLength 200 and -tbp 1. Tag directories combined both biological replicates per genotype. Aggregation plots were produced using ‘annotatePeaks.pl’ with -hist 10 and normalization set to the lowest sample tag count within a comparison group. Plots and corresponding bar charts, reported as tags per bp per peak, were visualized in GraphPad Prism 9.

Diffbind (v3.13) was used to generate a RangedSummarizedExperiment from BAM and narrowPeak files with the following parameters: fragmentLength=200, score=DBA_SCORE_READS, summits=TRUE, bUseSummarizeOverlaps=FALSE, bRemoveDuplicates=TRUE, and minOverlap=2 (Ross-Innes et al., 2012; Stark and Brown, 2011). mm10 ENCODE blacklist regions were removed by calling dba.blacklist(). Default normalization with full library size was called with dba.normalize(). ATAC-seq normalization included background=TRUE due to large changes between genotypes. RangedSummarizedExperiment was passed to DESeq2 for differential peak analysis using betaPrior=TRUE. Significance cutoff for differential peak analysis was considered fold change ≥ |2|, padj < 0.05. ATAC-seq MA plots were generated in Diffbind. PCA plots were generated using pcaExlorer on DESeq2 rlog transformed read counts.

### Enhancer Analysis

All genome arithmetic to compare peak coordinates was done using bedtools (Quinlan, 2014; Quinlan and Hall, 2010). This study only focused on active enhancer elements that were detected in one or more genotypes; regions enriched for H3K4me1 alone were excluded. Active enhancers were defined as a region with H3K4me1 and H3K27ac co-occupancy excluding intervals located within 2kb of a transcription start site or termination site (Calo and Wysocka, 2013; Creyghton et al., 2010). Chromatin accessibility was not used to define active enhancers due to the severe global depletion in ATAC-seq tag density observed in Pparα-null animals.

Gene coordinates were obtained from the UCSC Table Browser refGene (2020-08-17 update) (Karolchik et al., 2004). H3K4me1 and H3K27ac overlap was determined with a minimum 50% H3K27ac interval overlap over an H3K4me1 peak due to account for differences in histone mark average peak length. Fasting liver Pparα ChIP-seq coordinates were obtained from GSE118788 (Supplementary File Pparα-C57-Fast-peaks) (Sommars et al., 2019).

Differential enhancer groups were defined using the H3K27ac ChIP-seq DESeq2 output from above with the significance cutoff set at fold change ≥ |2|, padj < 0.05. DESeq2 normalized values were used for enhancer tag counts. Enhancer heatmaps were generated from bigWig files using deepTools 2.0 with ‘computeMatrix’ reference-point mode and ‘plotHeatmap’ (Ramírez et al., 2016).

Enhancer ontology was determined with the Genomic Regions Enrichment of Annotations Tool (GREAT) (McLean et al., 2010). GREAT version 4.0.4 was used in “basal plus extension” mode with default parameters and a 500kb maximum extension. Curated regulatory domains were included in calculations.

### Statistical Analysis

All statistical comparisons were carried out in GraphPad Prism 9 unless otherwise noted. Significance was determined using Student’s t-test, 1-way ANOVA with Tukey’s post hoc correction, or Kruskal-Wallis test with Dunn’s post hoc correction as noted. Shapiro-Wilk test for normality was used in R to determine whether to use a parametric or non-parametric test for significance for genomic data.

## DATA AND SOFTWARE AVAILABILITY

RNA-seq data were deposited in GSE165701. ChIP-seq and ATAC-seq data were deposited in GEO SuperSeries GSE179053. Scripts used for RNA-seq, ChIP-seq, and ATAC-seq analysis can be found at https://github.com/WolfgangLabJHMI/cpt2_ppara_liver_seq. The mass spectrometry proteomics data have been deposited to the ProteomeXchange Consortium via the PRIDE partner repository with the dataset identifier PXD027235.

## ACKNOWLEDGEMENTS

This work was supported in part by a National Institutes of Health grant R01DK120530 and R01DK116746 to M.J.W.

We thank Dr. Caitlyn Bowman-Cornelius for initial mouse breeding, Dr. Ebru Selen for advice on metabolomics analysis, and Dr. Anastasia Kralli for helpful conversations. We also thank Dr. Chan-Hyun Na and the Johns Hopkins Center for Proteomics Discovery for proteomic data acquisition.

## AUTHOR CONTRIBUTIONS

M.J.W and K.S.C conceptualized the project. K.S.C collected and analyzed data. K.S.C and M.J.W participated in the writing, review, and editing of the manuscript.

## Notes

### Competing Interest Statement

The authors have declared no competing interest.

