## Supplemental Figures for "Pparα and fatty acid oxidation coordinate hepatic transcriptional architecture"

#### SUPPLEMENTAL TABLES

Supplemental Table 1: RNA-seq differential expression analysis

Supplemental Table 2: Total proteome results

Supplemental Table 3: Acetyl-proteome results

Supplemental Table 4: Unbiased serum metabolomics

Supplemental Table 5: RNA-seq clusters gene list

Supplemental Table 6: Genome coordinates of active enhancers

#### SUPPLEMENTAL FIGURES

##### Supplement to Figure 1

A. Gene ontology for genes upregulated in $Cpt2^{L-/-}$ animals as determined by RNA-seq.
B. Gene ontology for peptides upregulated in $Cpt2^{L-/-}$ animals as determined by TMT-based quantitative mass spectrometry.
C. Volcano plot showing magnitude and significance for fasted liver proteome, measured by  (n=5). Peptide cell compartments are depicted.
D. Volcano plot showing magnitude and significance for fasted liver acetyl-proteome, measured by TMT-based quantitative mass spectrometry (n=5). Peptide cell compartments are depicted.
E. Acetyl-proteomics show Mitocarta peptides exhibit drastic hypoacetylation in $Cpt2^{L-/-}$ liver following a fast. Depicted are $\log_2$ WT/$Cpt2^{L-/-}$ fold changes for protein abundance (black) and acetyl-peptides. Acetyl-peptides are classified as hypoacetylated (red), no change (grey), or hyperacetylated (green). Fold change significance cutoff ≥ |2|, $p < 0.05$.
F. Genome browser tracks for representative genes that were either downregulated or showed no change in $Cpt2^{L-/-}$ RNA-seq. H3K9ac ChIP-seq signal trends with RNA-seq data (n=1).

##### Supplement to Figure 2

A. ATAC-seq genome browser tracks for representative genes that were downregulated or exhibited no change in $Cpt2^{L-/-}$ RNA-seq.

**Supplement to Figure 3**

A. PCA of untargeted serum metabolomics on fasted WT, Cpt2$^{L-/-}$, Ppar$\alpha^{-/-}$, and DKO liver (n=6).
B. Serum levels of free carnitine and short-, medium-, long-, and very long-chain acylcarnitines from fasted animals determined by median-scaled untargeted metabolomics (n=6, mean ± SEM)
C. Venn diagram depicting overlap between differentially expressed transcripts in Cpt2$^{L-/-}$, Ppar$\alpha^{-/}$, and DKO liver compared to WT as determined by RNA-seq.

RNA-seq significance cutoff is fold change ≥ |2|, padj < 0.05. One-way ANOVA followed by Tukey's post-hoc test was performed as appropriate. *p < 0.05; **p < 0.01; ***p < 0.001; ****p < 0.0001; ns, not significant.

**Supplement to Figure 4**

A. Heatmap showing all k-means clusters for differentially expressed RNA-seq transcripts. Transcript read counts are presented as row-scaled (Z-score) values.

**Supplement to Figure 5**

A. Aggregation plot depicting liver ATAC-seq mean tag density over all peak centers with ±5kb flanking regions.
B. PCA analysis of ATAC-seq from fasted WT, Cpt2$^{L-/-}$, Ppar$\alpha^{-/-}$, and DKO liver (n=2).

**Supplement to Figure 6**

A. Aggregation plot depicting liver H3K27ac ChIP-seq mean tag density over all peak centers with ±5kb flanking regions.

**Supplement to Figure 7**

A. Aggregation plot for ATAC-seq tag density over all active enhancers.
B. Aggregation plot for ATAC-seq tag density within 1kb of Pparα ChIP-seq binding sites within active enhancers.
C. Aggregation plots for H3K4me1 and H3K27ac tag density within 1kb of Pparα ChIP peak center at enhancers with increased H3K27ac signal in Cpt2KO liver.
D. (*top*) Pie chart showing genomic distribution of H3K4me1 differential peak analysis for enhancers with increased, repressed, and unchanged H3K27ac signal in DKO liver

compared to Cpt2KO. Promoters were not considered for H3K4me1 ChIP-seq. (*bottom*) Percent overlap between enhancers and Pparα ChIP-seq coordinates.

E. Bar graphs quantifying normalized H3K27ac and H3K4me1 ChIP-seq signal at Class C active enhancers.
F. Genome browser tracks for enhancer elements near *Pdk4* and *Fgf21*. Shown are ATAC-seq (red), H3K27ac ChIP-seq (purple), and H3K4me1 ChIP-seq (green). Active enhancer interval indicated by blue bar, Ppara ChIP-seq binding site indicated by black bar. TSS = transcription start site, TTS = transcription termination site.

Significance determined by Kruskal-Wallis test with Dunn's post hoc correction. *p < 0.0001; ns, not significant. Bar graphs represent 2%-98% percentile.

### Fig S1

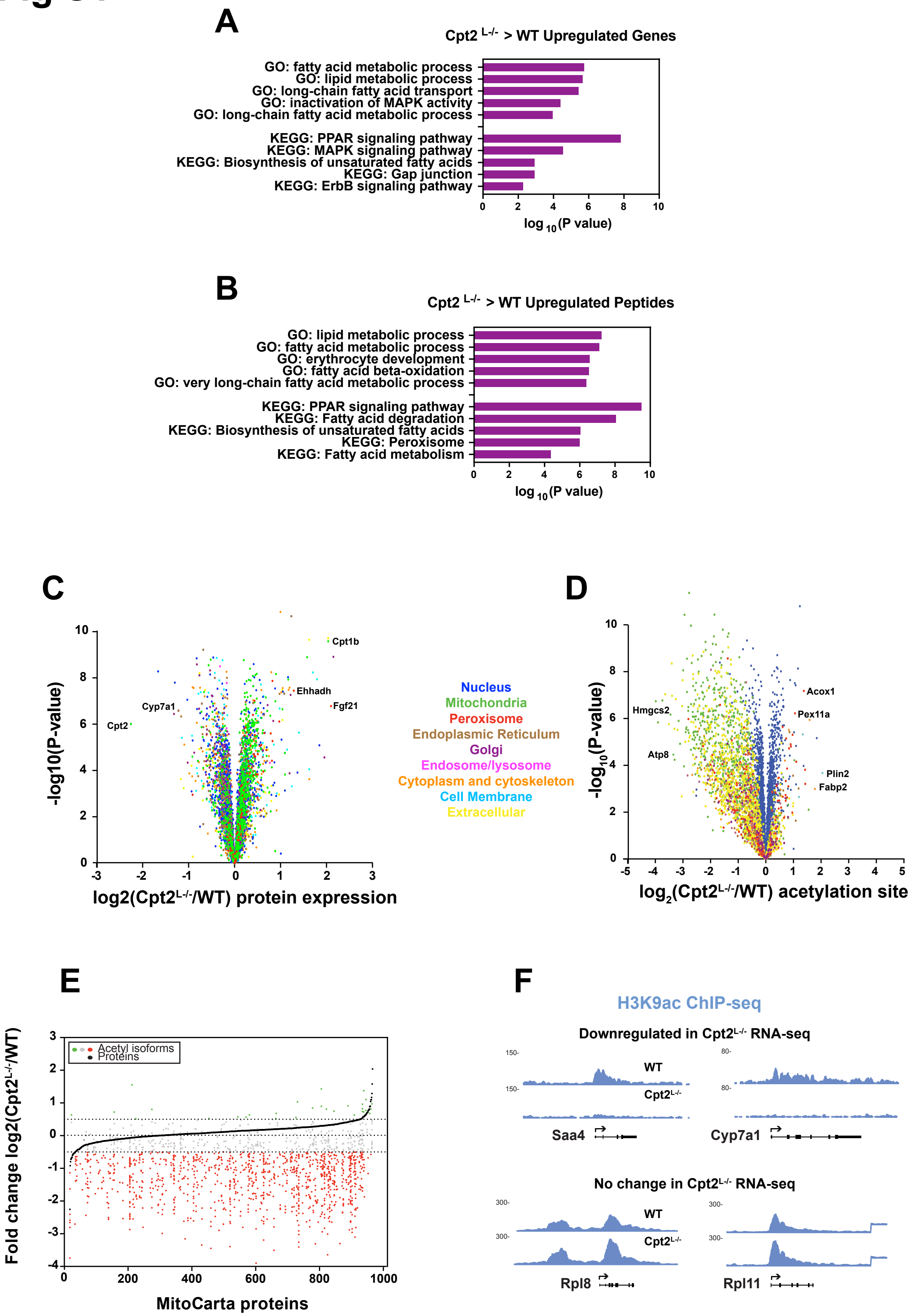
A
Cpt2 L-/- > WT Upregulated Genes
GO: fatty acid metabolic process
GO: lipid metabolic process
GO: long-chain fatty acid transport
GO: inactivation of MAPK activity
GO: long-chain fatty acid metabolic process
KEGG: PPAR signaling pathway
KEGG: MAPK signaling pathway
KEGG: Biosynthesis of unsaturated fatty acids
KEGG: Gap junction
KEGG: ErbB signaling pathway
log 10 (P value)
B
Cpt2 L-/- > WT Upregulated Peptides
GO: lipid metabolic process
GO: fatty acid metabolic process
GO: erythrocyte development
GO: fatty acid beta-oxidation
GO: very long-chain fatty acid metabolic process
KEGG: PPAR signaling pathway
KEGG: Fatty acid degradation
KEGG: Biosynthesis of unsaturated fatty acids
KEGG: Peroxisome
KEGG: Fatty acid metabolism
log 10 (P value)
C
-log10(P-value)
Cpt1b
Ehhadh
Fgf21
Cyp7a1
Cpt2
log2(Cpt2L-/-/WT) protein expression
Nucleus
Mitochondria
Peroxisome
Endoplasmic Reticulum
Golgi
Endosome/lysosome
Cytoplasm and cytoskeleton
Cell Membrane
Extracellular
D
-log10(P-value)
Hmgcs2
Atp8
Acox1
Pex11a
Plin2
Fabp2
log2(Cpt2L-/-/WT) acetylation site
E
Fold change log2(Cpt2L-/-/WT)
Acetyl isoforms
Proteins
MitoCarta proteins
F
H3K9ac ChIP-seq
Downregulated in Cpt2L-/- RNA-seq
WT
Cpt2L-/-
Saa4
Cyp7a1
No change in Cpt2L-/- RNA-seq
WT
Cpt2L-/-
Rpl8
Rpl11

### Fig S2

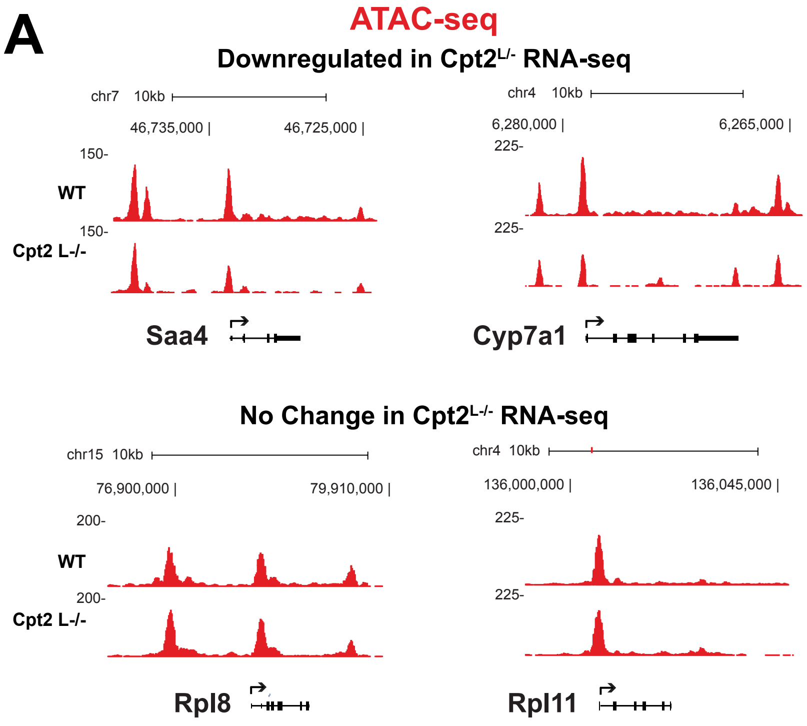
A
ATAC-seq
Downregulated in Cpt2L/- RNA-seq
chr7 10kb
46,735,000 |
46,725,000 |
150-
WT
150-
Cpt2 L-/-
Saa4
chr4 10kb
6,280,000 |
6,265,000 |
225-
225-
Cyp7a1
No Change in Cpt2L-/- RNA-seq
chr15 10kb
76,900,000 |
79,910,000 |
200-
WT
200-
Cpt2 L-/-
Rpl8
chr4 10kb
136,000,000 |
136,045,000 |
225-
225-
Rpl11

### Fig S3

A

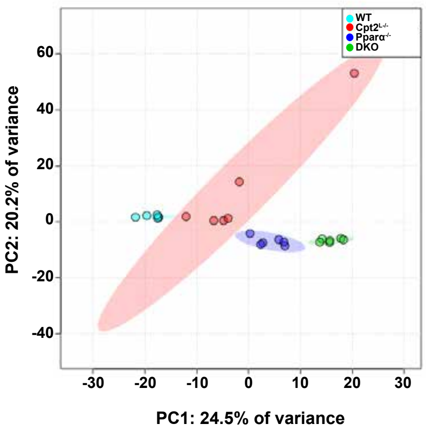


B

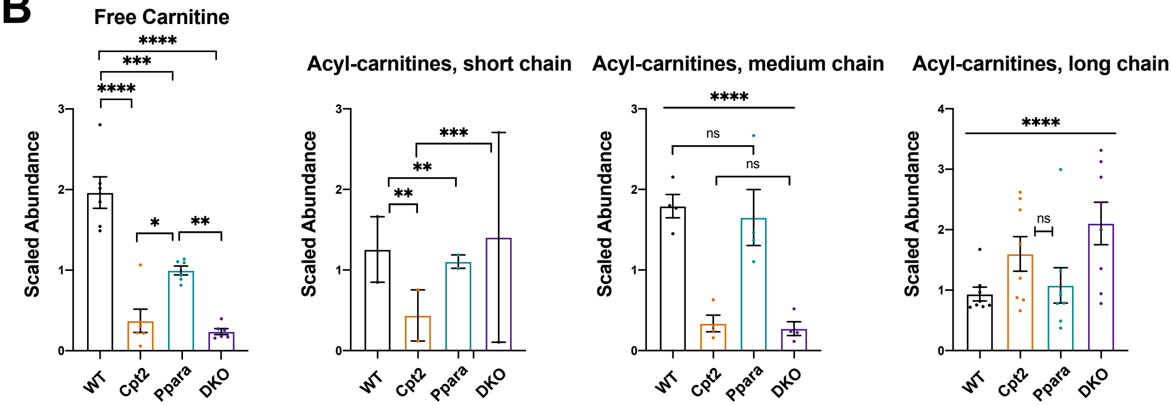


C

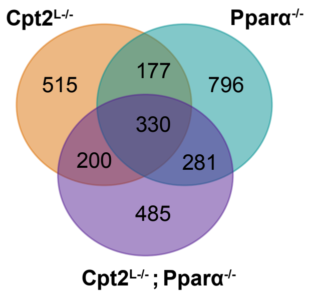


All differentially expressed genes (KO/WT)
fold change ≥ ±2, padj < 0.05

### Fig S4

A

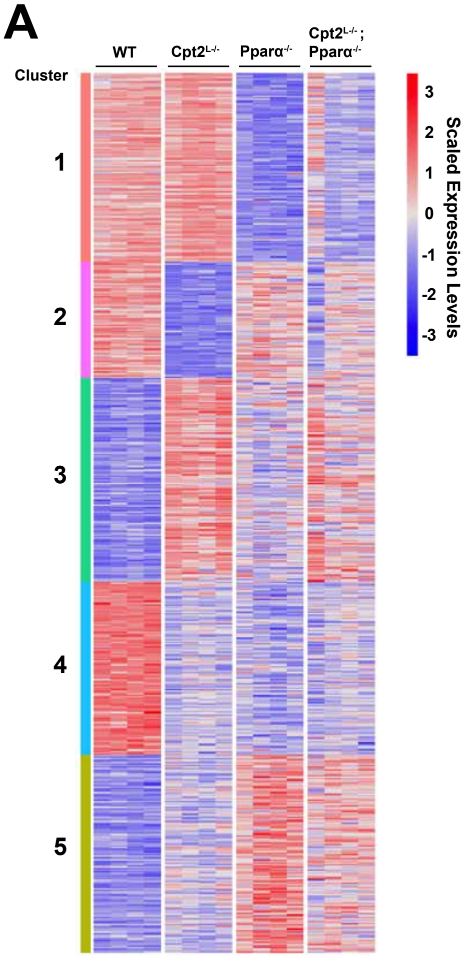
WT
Cpt2L-/-
Pparα-/-
Cpt2L-/-; Pparα-/-
Cluster
1
2
3
4
5
3
2
1
0
-1
-2
-3
Scaled Expression Levels

### Fig S5

A

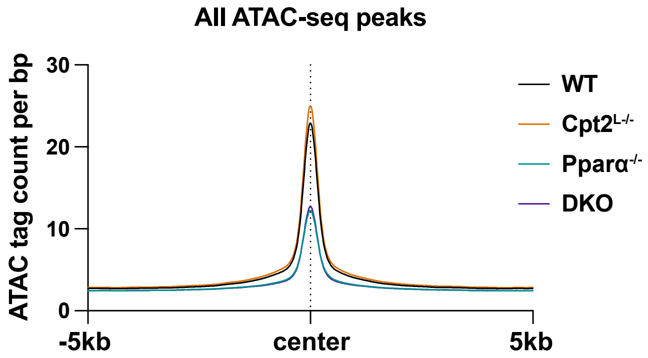
All ATAC-seq peaks
ATAC tag count per bp
30
20
10
0
-5kb
center
5kb
WT
Cpt2L-/-
Pparα-/-
DKO

B

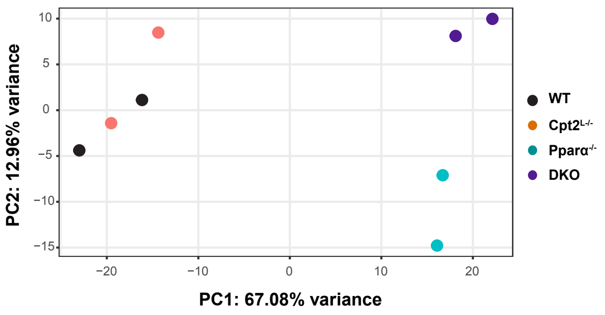
PC2: 12.96% variance
PC1: 67.08% variance
10
5
0
−5
−10
−15
−20
−10
0
10
20
WT
Cpt2L-/-
Pparα-/-
DKO

### Fig S6

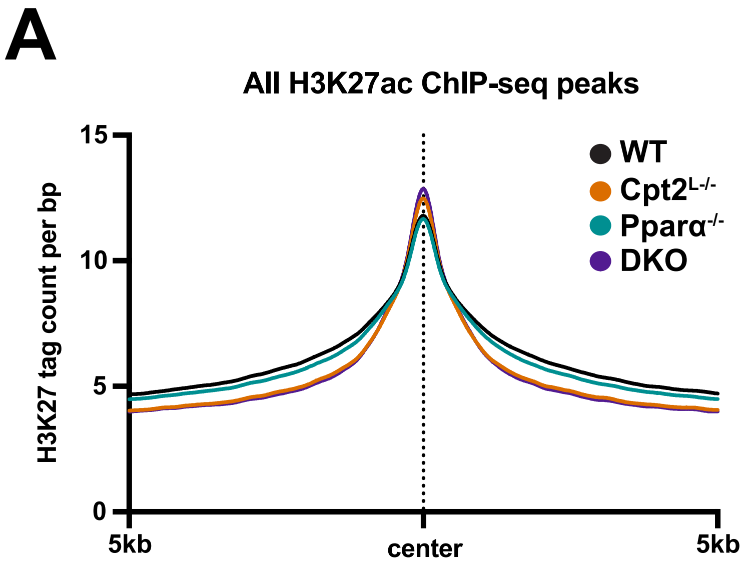
A
All H3K27ac ChIP-seq peaks
H3K27 tag count per bp
15
10
5
0
5kb
center
5kb
WT
Cpt2L-/-
Pparα-/-
DKO

### Fig S7

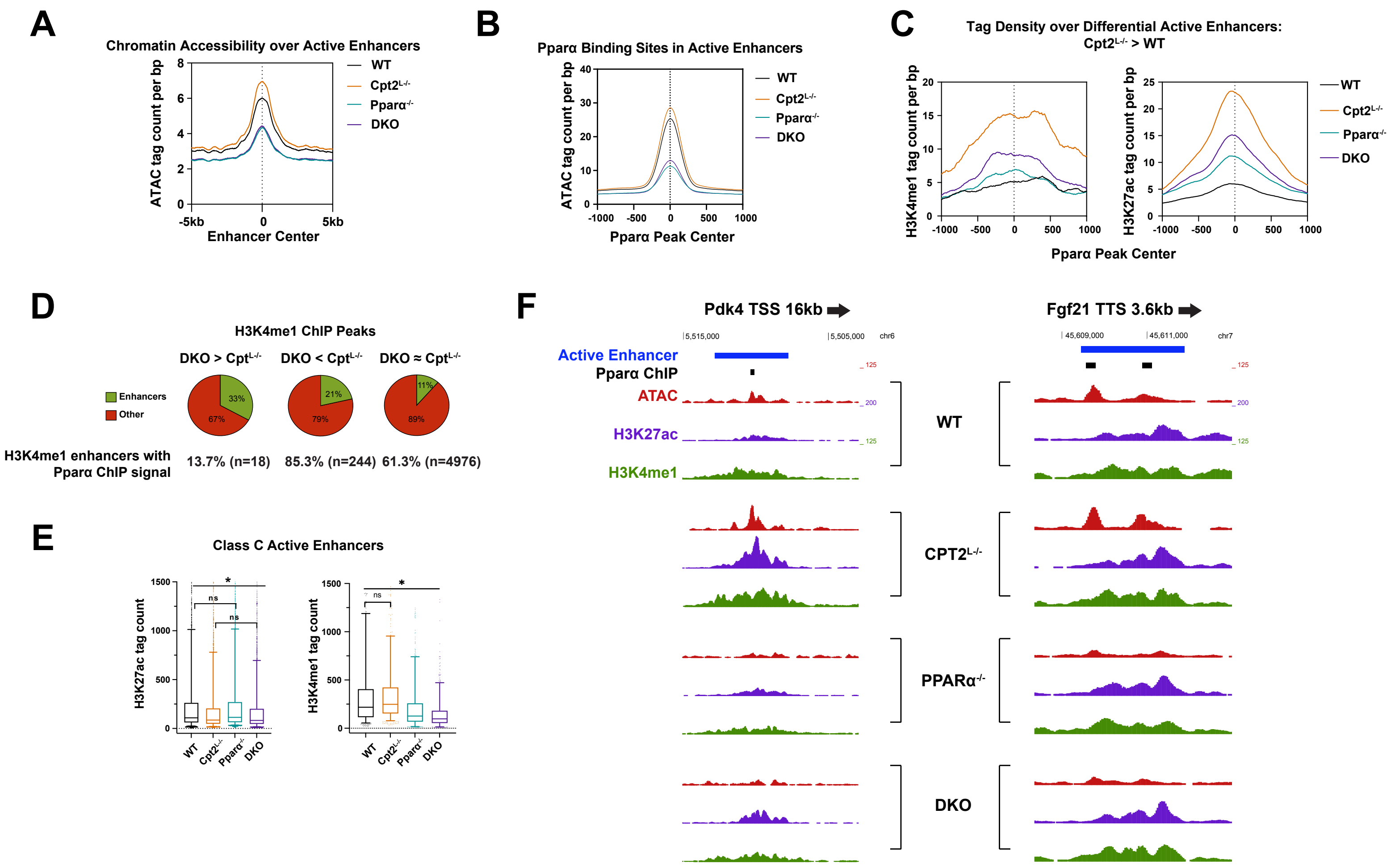
A
Chromatin Accessibility over Active Enhancers
ATAC tag count per bp
Enhancer Center
-5kb
0
5kb
WT
Cpt2L-/-
Pparα-/-
DKO
B
Pparα Binding Sites in Active Enhancers
ATAC tag count per bp
Pparα Peak Center
C
Tag Density over Differential Active Enhancers:
Cpt2L-/- > WT
H3K4me1 tag count per bp
H3K27ac tag count per bp
Pparα Peak Center
D
H3K4me1 ChIP Peaks
DKO > CptL-/-
DKO < CptL-/-
DKO ≈ CptL-/-
Enhancers
Other
33%
67%
21%
79%
11%
89%
H3K4me1 enhancers with
Pparα ChIP signal
13.7% (n=18)
85.3% (n=244)
61.3% (n=4976)
E
Class C Active Enhancers
H3K27ac tag count
H3K4me1 tag count
ns
*
WT
Cpt2L-/-
Pparα-/-
DKO
F
Pdk4 TSS 16kb
Fgf21 TTS 3.6kb
5,515,000
5,505,000
chr6
45,609,000
45,611,000
chr7
Active Enhancer
Pparα ChIP
ATAC
H3K27ac
H3K4me1
WT
CPT2L-/-
PPARα-/-
DKO
